# Extrahepatic, cell-specific delivery of LNPs through competitive inhibition of ApoE-mediated uptake

**DOI:** 10.64898/2026.05.29.727812

**Authors:** Joshua C. Bufton, Thomas I.P. Green, Angharad Walters, Adam W. Perriman, Benjamin M. Carter

**Affiliations:** Hone Bio Limited, Science Creates Old Market, Midland Road, Bristol, BS2 0JZ; School of Cellular and Molecular Medicine, University of Bristol, University Walk, Bristol, BS8 1TD UK; Research School of Chemistry, Australian National University, Building 137 Sullivans Creek Rd, Acton, Canberra, ACT 2601 Australia; John Curtin School of Medical Research, Australian National University, 131 Garran Rd, Acton, Canberra, ACT 2601 Australia; ARC Centre of Excellence in Synthetic Biology, Australian National University, Canberra, ACT 2601, Australia

## Abstract

Targeted lipid nanoparticles (LNPs) for extrahepatic drug delivery are limited by apolipoprotein E (ApoE)-mediated hepatic accumulation. We developed NanoPilot^™^, a modular fusion protein platform comprising antibodies and anchors blocking the low density lipoprotein receptor (LDLR) ApoE binding site, to block LNP liver uptake and redirect to target cells. NanoPilot can be applied to preformulated LNPs in 10 minutes with two pipetting steps.

*In vitro*, an anti-CD3ε NanoPilot increased T-cell transfection 40-fold and reduced monocyte transfection 10-fold in human peripheral blood mononucleocytes. In immunocompetent mouse models, NanoPilot-coated LNPs achieved 30–40% splenic and hepatic T-cell transfection whilst bulk liver accumulation was reduced 3-fold. An anti-c-Kit NanoPilot was further shown to enhance delivery to a haematopoietic stem cell-like cell line in an *in-vitro* co-culture assay.

NanoPilot establishes a versatile framework for the systemic delivery of genetic therapies through concomitant cell-specific targeting and off-target blocking. Further research will assess potential clinical applications.

## Introduction and background

Lipid nanoparticles (LNPs) are a drug-delivery platform comprising a lipid-based membrane, a therapeutic cargo and optional targeting ligands.^1^ Upon administration, LNPs acquire a protein corona that, together with any targeting ligands, influences their destination.^2, 3^ Ideally, LNPs bind to a cell membrane receptor on the target cell and are endocytosed, releasing their payload into the cytoplasm and completing transfection.^2^ In reality, some LNPs do not reach the target cells or fail to complete endosomal escape of their payload.^4, 5^ Moreover, the magnitude of translation of delivered mRNA correlates weakly with the quantity of mRNA delivered, and translational efficiency is affected by the targeting ligand used.^6^

LNP technology has a history going back more than six decades.^7^ Recently, focus has turned to the potential that LNPs hold for delivering gene therapies, mRNA-based cancer vaccines and cell therapies.^7–12^ The recent surge in LNP clinical translation is driven by their scalable manufacturing, ability to encapsulate diverse and large genetic payloads, and reduced immunogenicity relative to viral platforms.^1, 7, 10^ Indeed, the utility of LNP technology has been proven in three approved drugs: two mRNA vaccines against COVID-19 and an siRNA therapeutic.^13–18^

Wider incorporation of LNPs into therapeutics has been limited by off-target delivery, which has consequences of poor target cell transfection efficiency and potential for adverse effects following delivery to inappropriate cell types.^5, 19^ In particular, LNPs tend to target hepatocytes via interaction with apolipoprotein E (ApoE), a component of the protein corona.^3, 5^ LNPs with bound ApoE then bind to low density lipoprotein receptors (LDLR) and are consequently taken into hepatocytes,^5^ although LDLR are near-ubiquitous.^20^ Hepatic delivery is useful for therapeutics targeting liver disease, such as patisiran (approved for the treatment of hereditary transthyretin-mediated amyloidosis,^14, 17^) but limits the wider application of LNPs. It was recently shown that hepatocyte expression blunts the effectiveness of cancer vaccines in mice, for example.^21^

Several approaches to improve LNP targeting by modifying the ApoE–LNP interaction have been reported. Park et al. and Theuerkauf et al. exploited apolipoproteins’ propensity to bind LNPs for anchoring antibodies and designed ankyrin repeat proteins (DARPins), respectively, to LNP surfaces.^22, 23^ Kayabolen et al. engineered a ‘dead’ ApoE (non-LDLR-binding) and used hyperactive PCSK9 to reduce ApoE–LDLR-mediated uptake and re-target with antibodies.^24^ Other groups have attempted to mitigate off-target effects by modifying the lipids in the LNP.^11, 25^ However, each approach has its limitations and off-target delivery is still observed.

The LNP with maximum suitability for drug delivery would demonstrate reduced-to-no off-target delivery and enhanced cell-specific targeting. Such an LNP could be used in cell and gene therapies, including *in-vivo* chimeric antigen receptor (CAR)-T cancer or autoimmune therapies without lymphodepletion,^1, 11, 12, 26, 27^ targeted gene therapy,^9^ and potentially curative HIV therapies.^28^

NanoPilot is an IgG-based biologic designed to control LNP fate through both redirect and blocking functions. It consists of an antibody scaffold with the ApoE-binding domains of LDLR fused to the C-terminus of each heavy chain, with the hypothesis that this would both reduce liver sequestration and enhance payload delivery to target cells. We report here the characterisation and optimisation of the NanoPilot platform and its first applications *in vitro* and *in vivo*.

### NanoPilot design and validation

NanoPilot is a fusion protein built on a human IgG1 backbone (**Fig. 1A**). LALA-PG mutations in the backbone silence Fc receptor binding,^29^ preventing non-specific uptake by immune cells and reducing antibody-dependent cellular cytotoxicity.^30^

**Figure 1.**
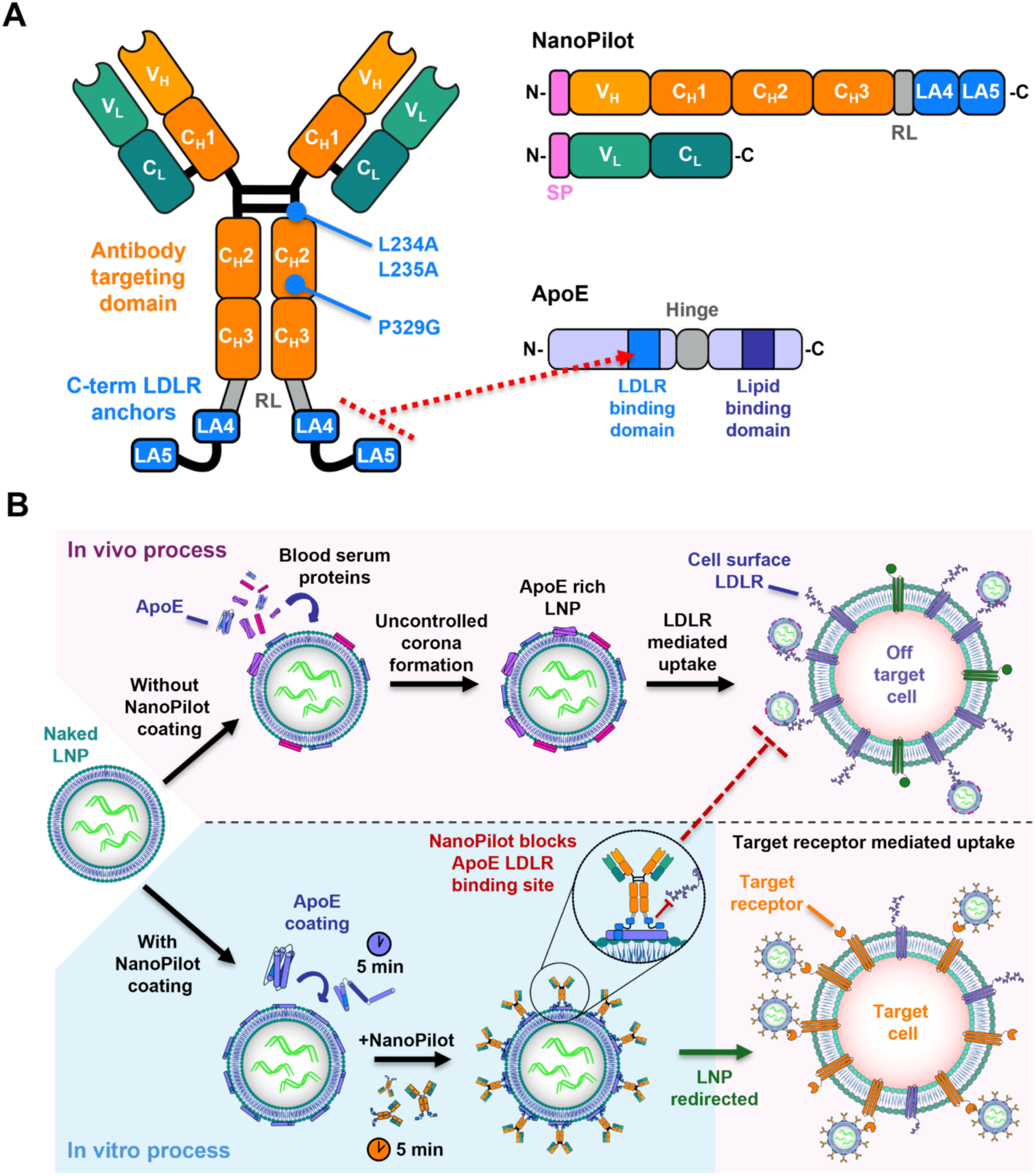
Overview of the novel NanoPilot technology. **(A)** Structure and domain organisation of the NanoPilot system, comprising antibodies for cell-specific targeting and anchors to block the LDLR binding site of ApoE. LALA-PG mutations to silence Fc interactions are shown. **(B)** Overview of LDLR-mediated uptake, the ∼10-minute NanoPilot functionalisation process, and the block-and-redirect mechanism. ApoE, apolipoprotein E; C_H_X, constant heavy region X; C_L_, constant light region; LA, LDLR type A domain; LDLR, low density lipoprotein receptor; RL, rigid linker; SP, secretion/signal peptide; V_H_, variable heavy chain; V_L_, variable light chain. We first utilised a patisiran-like LNP formulation (MC3-DSPC) to evaluate NanoPilot, given its well-established therapeutic use for liver cell delivery via ApoE–LDLR endocytosis.^14, 17^. Dynamic light scattering (DLS) verified the homogeneity and structural integrity of the assembled LNPs **(Supplementary Table 1; Supplementary** Fig. 2A).

Variable regions from well-characterised antibodies (e.g. OKT3) bind with high affinity to target receptors and provide the redirect function of NanoPilot (**Supplementary Fig. 1A–D)**. ApoE blocking is achieved by fusing ApoE-binding regions of human LDLR (LA4–LA5 domains) to the protein (**Fig. 1A, B**). ApoE comprises a C-terminal lipid-binding domain and an N-terminal receptor-binding domain, which are connected by a flexible hinge region (**Fig. 1A**). Crucially, the N-terminal motif responsible for LDLR engagement remains sterically shielded and is only exposed following a conformational change upon binding to lipoproteins or LNPs.^31^ Therefore, NanoPilot only binds ApoE in its membrane-bound conformation and not free ApoE in solution (**Fig. 1B**). The LDLR fragment is attached to the C-terminus of each IgG heavy chain, which ensures the variable domains are oriented away from the LNP surface. This bivalent display is designed to increase the avidity to the multiple ApoE proteins present on the LNP surface.

NanoPilot is applied to existing LNPs carrying a payload, in a simple two-step non-covalent functionalisation process that takes 10 minutes and is amenable to automation and high-throughput screening (**Fig. 1B**). Size exclusion and SDS-PAGE assays of purified NanoPilot material showed homogeneity and lack of aggregates, and demonstrated the antibody is intact, folded and held together via disulfide bonds (**Supplementary Fig. 1E, F**). Indirect ELISAs of the purified NanoPilot molecules show that high affinity antigen binding remains intact (**Supplementary Fig. 1A–D**).

To evaluate the block and redirect mechanism of an anti-human CD3ε NanoPilot (hCD3-NP), we used two LDLR-expressing Jurkat T cell lines: one positive and one negative for a CD3ε target antigen (CD3+ and CD3-, respectively). We confirmed that the addition of ApoE3 to non-NanoPilot LNPs led to equal transfection of both CD3+ and CD3-cell lines (**Fig. 2A**), establishing the suitability of the model for assessing the block and redirect mechanism of hCD3-NP.

**Figure 2.**
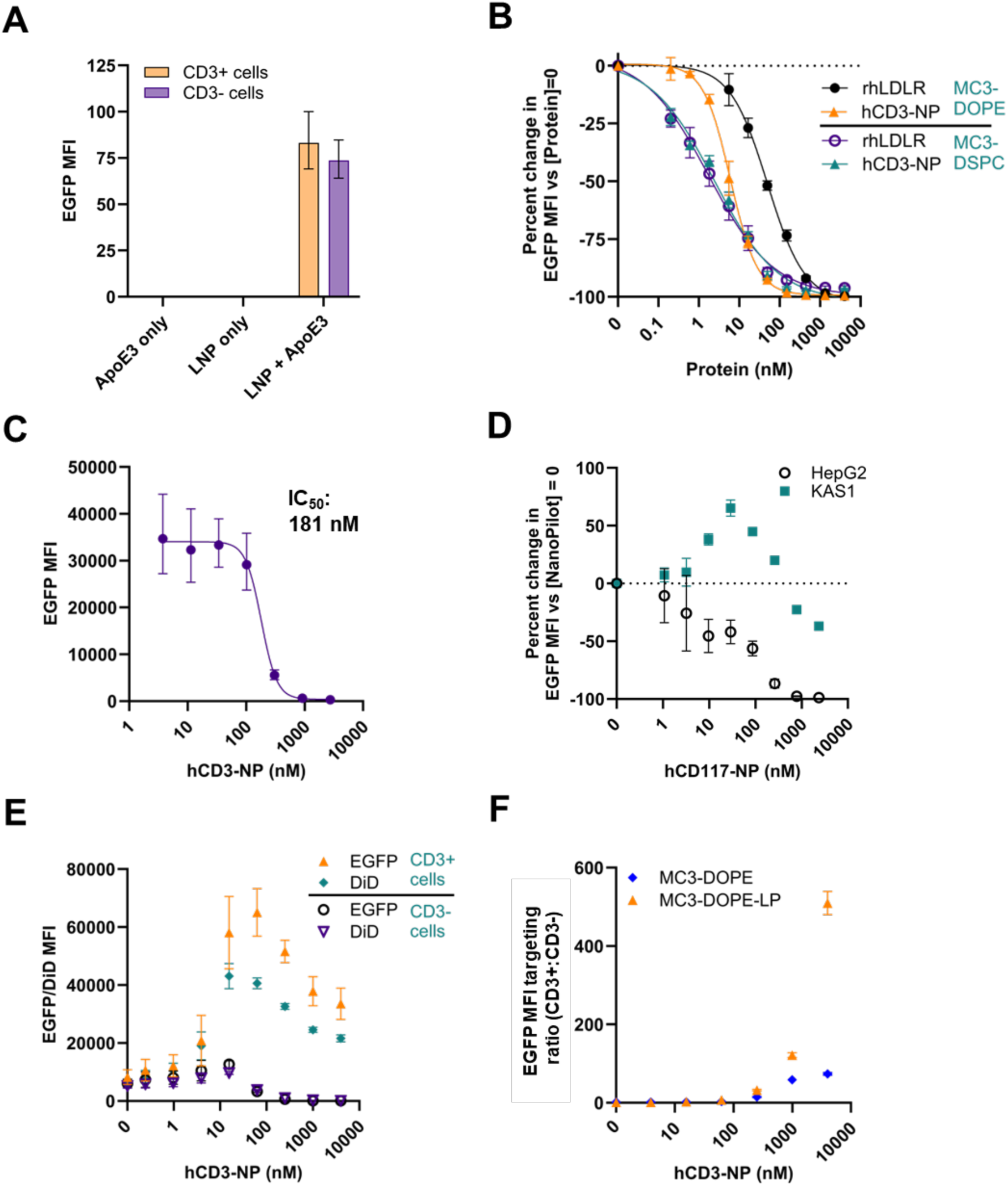
NanoPilot inhibits non-specific ApoE3-mediated uptake and redirects EGFP mRNA delivery to target human cell lines *in vitro*. **(A)** Flow cytometry analysis of EGFP expression in a co-culture of CD3+ and CD3-Jurkat T cells after treatment with MC3-DSPC LNP ± ApoE3. **(B)** Dose– response curve in a monoculture of CD3-Jurkat cells with MC3-DSPC and MC3-DOPE LNPs, showing inhibition of ApoE3–LNP-mediated EGFP expression by the hCD3-NP (OKT3 clone) and rhLDLR. IC_50_ values are listed in **Supplementary Table 2**. **(C)** Dose–response curve showing inhibition of MC3-DSPC LNP-mediated EGFP expression by hCD3-NP in HepG2 monoculture. X-axis values and determined IC_50_ value represents the concentration of hCD3-NP used for LNP coating. See **Supplementary Note** for specific methods. **(D)** EGFP expression in a co-culture of HepG2 cells and CD117+ Kasumi-1 cells, treated with a titration of hCD117-NP (9P3 clone) with MC3-DSPC LNP. See **Supplementary Note** for specific methods. **(E)** MC3-DOPE-LP transfection of CD3-/CD3+ co-culture. Uptake (DiD MFI) and mRNA translation (EGFP MFI) correlate in target and off-target cells. **(F)** Targeting ratio for MC3-DOPE-LP LNPs, showing enhanced on-target delivery and minimal off-target uptake with increasing concentrations of hCD3-NP versus MC3-DOPE LNP. Data are presented as mean ± SD (n=3). ApoE3, apolipoprotein E3; EGFP, enhanced green fluorescent protein; hCD117, human CD117; hCD3, human CD3; IC_50_, half maximal inhibitory concentration; KAS1, Kasumi-1 cells; LNP, lipid nanoparticle; MFI, median fluorescence intensity; NP, NanoPilot; PBS, phosphate-buffered saline; rhLDLR, recombinant human low density lipoprotein receptor; SD, standard deviation.

We assessed the capacity of hCD3-NP to block ApoE3-mediated uptake of MC3-DSPC LNPs in both CD3- and HepG2 (a human hepatocellular carcinoma cell line) monocultures, recording IC_50_ values of 2.4 nM and 181 nM, respectively (**Fig. 2B, C; Supplementary Table 2**). The IC_50_ values reflect the concentration of NanoPilot added to the LNPs during the initial coating step, rather than the final diluted concentration in the cell culture media.

Next we evaluated the NanoPilot block and redirect mechanism using the CD3+/CD3-co-culture model and two anti-CD3 NanoPilot clones (derived from OKT3 and cibisatamab variable sequences). Flow cytometry revealed that the OKT3-NanoPilot led to higher EGFP expression in target CD3+ cells compared to the cibisatamab-derived version (**Supplementary Fig. 3A, B**). Addition of the hCD3-NP to MC3-DSPC LNPs increased EGFP expression by target CD3+ cells compared with the bare LNP (**Supplementary Fig. 3A**). To quantify specificity, we calculated targeting ratios (target/off-target fluorescence intensity); the cibisatamab- and OKT3-derived NanoPilots achieved 25- and 40-fold ratios, respectively (**Supplementary Fig. 3B**). Later experiments therefore used the OKT3 clone. Similar block and redirect effects were observed with an anti-c-KIT NanoPilot (hCD117-NP) coating MC3-DSPC LNPs applied to a co-culture of HepG2 and Kasumi-1 cells (**Fig. 2D**). Transfection of target-positive cells is biphasic with respect to NanoPilot concentration: initially increasing above baseline, with signal dropping off at higher concentrations (although the targeting ratio peaks later); Hashiba et al. determined the similar bell-shaped response of single-domain antibody (VHH)-modified LNPs to be due to receptor degradation.^32^

To further optimise CD3+ Jurkat targeting, we screened variants of the MC3-DSPC formulation. Substituting the DSPC helper lipid with DOPE (MC3-DOPE) increased target cell transfection and improved the targeting ratio to >200-fold **(Supplementary Fig. 4A, B)**. While hCD3-NP and the LDLR extracellular domain (rhLDLR) exhibited comparable IC_50_ values for MC3-DSPC LNPs, hCD3-NP achieved an ∼8-fold lower IC_50_ than rhLDLR for MC3-DOPE LNPs (6.04 nM vs 49.33 nM, respectively; **Fig. 2B; Supplementary Table 2)**. Reducing the DMG-PEG2000 content in the MC3-DOPE LNP formulation to 0.75% (MC3-DOPE-LP) further boosted the targeting ratio to >500-fold **(Fig. 2E, F; Supplementary Fig. 4C)**.

To verify that this specificity was driven by uptake rather than post-entry translation, we incorporated a C18-anchored DiD dye to track LNPs independently of mRNA expression. The strong correlation between DiD signal and EGFP expression (r=0.98 for CD3+ and r=0.99 for CD3-cells; P<0.0001) indicates that NanoPilot acts primarily at the level of cellular uptake, physically restricting LNP entry into off-target cells while directing it toward the target population **(Fig. 2E)**. DLS analyses of the original and optimised formulations show negligible size differences, including after addition of ApoE3 **(Supplementary Table 1; Supplementary Fig. 2A, B)**.

We next sought to confirm that the enhanced uptake in CD3+ cells was driven by NanoPilot– LNP engagement rather than OKT3-mediated CD3ε stimulation. To test this, we evaluated MC3-DOPE-LP LNPs formulated with either hCD3-NP alone (without ApoE3) or an OKT3 control antibody in a CD3+/CD3-co-culture system **(Supplementary Fig. 4D, E)**. Neither hCD3-NP alone nor OKT3 combined with ApoE3 yielded targeted cellular uptake, demonstrating that CD3+ targeting specifically requires the co-presence of hCD3-NP and ApoE3.

Time-resolved microscopy showed an increased rate of EGFP expression for CD3+ cells with hCD3-NP concentrations ≥62.5 nM, while the rate of CD3-uptake was reduced with hCD3-NP concentrations ≥250 nM **(Fig. 3A, B)**. Area under the curve analysis of the time-resolved data was in agreement with the FACS data **( Fig. 3C)**.

**Figure 3.**
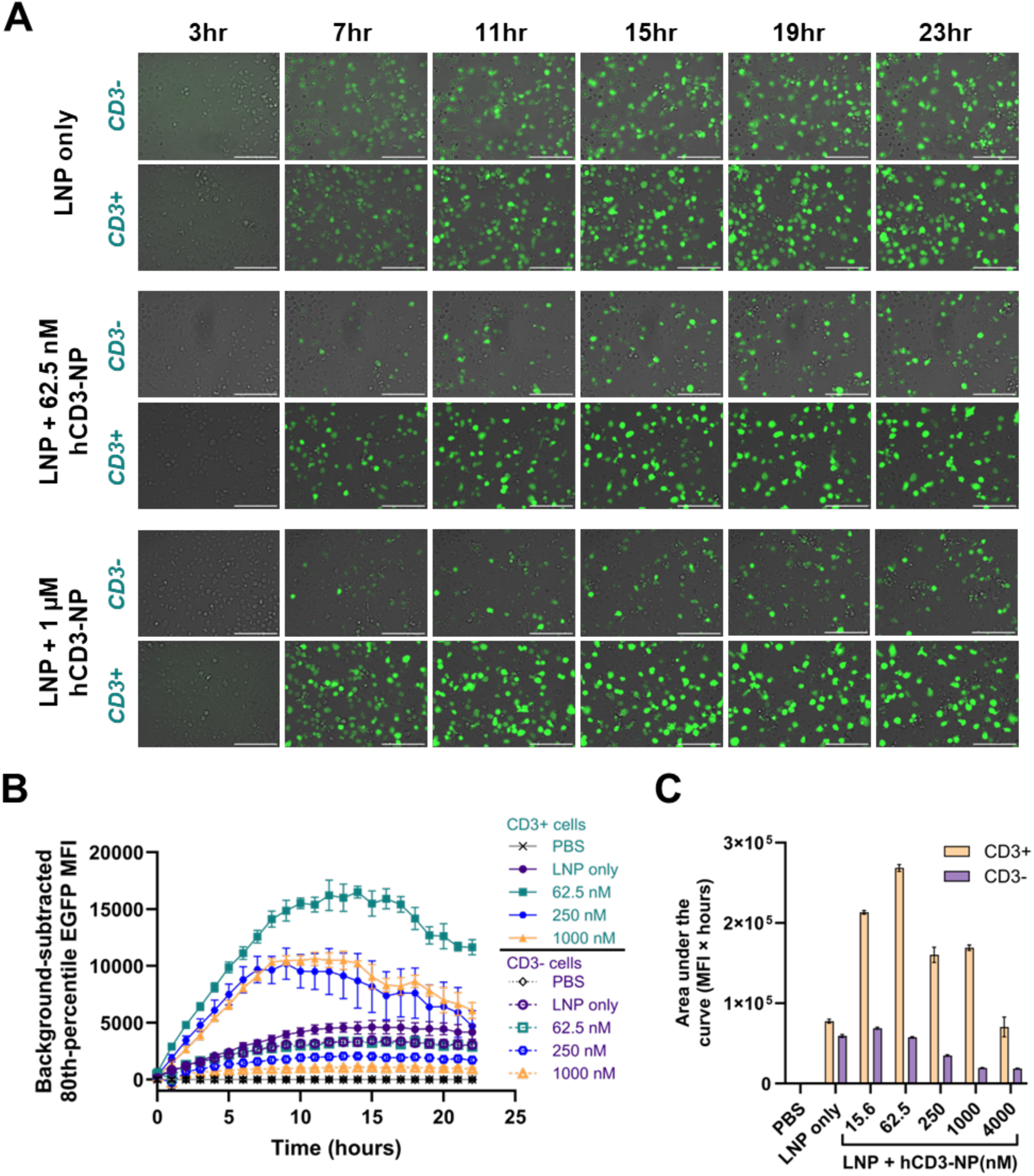
Time-resolved microscopy analysis of transfection. **(A)** Images of CD3- and CD3+ Jurkat monocultures treated with MC3-DOPE-LP LNPs ± hCD3-NP at low (62.5 nM) and high (1 µM) coating concentrations. Jurkat T cell imaging methods can be found in the **Supplementary Note**. **(B)** Background-subtracted EGFP MFI values plotted for each condition across all imaged timepoints. Data are presented as mean ± SD (n=3). **(C)** Corresponding area under the curve analysis of plotted time course showing increased on-target and reduced off-target EGFP expression with hCD3-NP versus uncoated LNP. Plotted bars represent the mean of three replicate wells ± standard error of the mean. EGFP, enhanced green fluorescent protein; hCD3, human CD3; LNP, lipid nanoparticle; MFI, median fluorescence intensity; NP, NanoPilot; PBS, phosphate-buffered saline.

### *In-vivo* targeting of murine T cells

Prior to *in-vivo* studies, we validated a murinised NanoPilot on murine cells *in vitro*. We initially titrated an anti-murine CD3ε NanoPilot (mCD3-NP) on MC3-DOPE-LP LNP concentrated to 100 μg/mL as required for *in-vivo* dosing. Compared with LNP alone, mCD3-NP showed increased transfection of isolated activated CD4+ splenocytes with an optimal coating concentration of 500 nM–1 µM (**Fig. 4A**). In unactivated murine splenocytes, mCD3-NP and an anti-murine CD8α NanoPilot (mCD8-NP) targeted LNPs to CD4+ and CD8+, and CD8+ cells, respectively (**Fig. 4B, C**). Fluorescence intensity data revealed a decoupling of LNP uptake and EGFP translation: despite mCD8-NP mediating higher LNP–cell interaction (DiD MFI) in CD8+ cells relative to mCD3-NP, EGFP expression was considerably lower (**Supplementary Fig. 5A, B**). CD4+ cell counts were comparable in LNP-only and LNP+NanoPilot conditions whereas CD8+ cell counts were decreased in the LNP+mCD8-NP condition (**Supplementary Fig. 5C**). Assessment of the late activation marker CD25 showed upregulation by the mCD3-NP–LNP complex, an effect absent with uncomplexed mCD3-NP, and mCD8-NP ± LNP groups **(Supplementary Fig. 5D)**.

**Figure 4.**
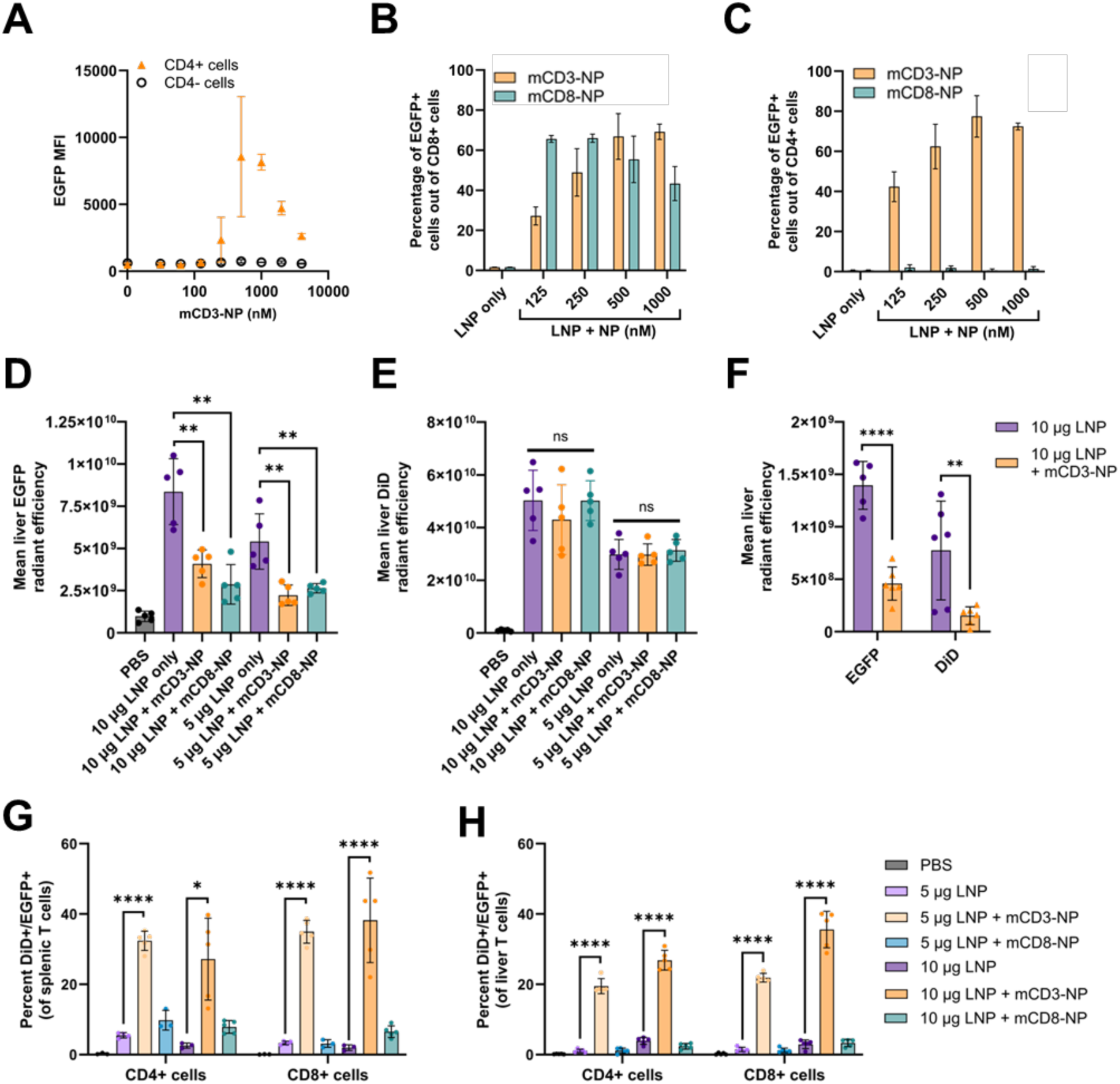
NanoPilot facilitates targeted LNP delivery to murine T cells *in vitro* and *in vivo*, while reducing hepatic accumulation versus controls. **(A)** Flow cytometric quantification of *in-vitro* EGFP expression in CD3/CD28 bead-activated primary murine CD4+ T cells (isolated from BALB/c splenocytes) treated with MC3-DOPE-LP LNPs formulated with a titration range of mCD3-NP. **(B)** Selective targeting in unactivated murine pan T cell cultures (splenocyte-derived) treated with LNPs ± mCD3-NP and mCD8-NP, showing transfection of CD8+ and **(C)** CD4+ T cells. (D, E) *Ex-vivo* IVIS imaging of C57BL/6 livers 24 hours post-injection. Quantification of the average radiant efficiency of the **(D)** EGFP reporter and **(E)** DiD lipid tracer demonstrates a significant reduction in hepatic EGFP translation when LNPs are coated with NanoPilot but no significant change in DiD signal. (F) *Ex-vivo* IVIS imaging of BALB/c livers 48 hours post-injection. Radiant efficiency quantification reveals a significant decrease in both LNP accumulation (DiD) and payload expression (EGFP) with mCD3-NP coating versus uncoated LNP. **(G)** Flow cytometric analysis of C57BL/6 splenocytes and **(H)** liver-resident T cells 24 hours post-injection, showing that mCD3-NP facilitated significant increases in targeted LNP uptake and EGFP expression compared with LNP alone. Data represent the percentage of DiD+/EGFP+ cells within the respective CD4+ and CD8+ T-cell populations. Data are presented as mean ± SD (n=5 biologically independent animals per group, except for the splenic PBS [n=3], splenic 10 µg untargeted LNP [n=4] and liver PBS [n=4] controls, due to sample loss during *ex-vivo* tissue processing). *P<0.05, **P<0.01, ***P<0.001, ****P<0.0001. EGFP, enhanced green fluorescent protein; IVIS, *in vivo* imaging system; LNP, lipid nanoparticle; mCD3/4/8, murine CD3/4/8; MFI, median fluorescence intensity; NP, NanoPilot; ns, not significant; PBS, phosphate-buffered saline; SD, standard deviation.

Next we sought to evaluate NanoPilot T-cell targeting and ApoE blocking *in vivo*. *In vivo* imaging system (IVIS) imaging of murine livers at 24 hours post-LNP injection revealed a divergence between LNP accumulation and EGFP expression **(Fig. 4D, E; Supplementary Fig. 6A)**. While liver DiD radiant efficiency remained relatively stable across groups, hepatic EGFP expression was significantly attenuated (∼3-fold; P<0.01) when LNPs were formulated with NanoPilot. This reduction in liver EGFP versus the LNP-only groups was observed for both the mCD3-NP and mCD8-NP groups **(Fig. 4D)**. In a separate murine study evaluating *ex-vivo* liver signal at 48 hours, we observed a significant, ∼3-fold decrease in both EGFP and DiD signals (P<0.0001 and P<0.01, respectively; **Fig. 4F**) when mCD3-NP was added to LNPs. This suggests that the sustained DiD levels measured at 24 hours likely reflect LNPs still in systemic circulation or transiently associated with the liver vasculature, which are then markedly reduced by the 48-hour mark.

At the cellular level, flow cytometric analysis of spleen- and liver-resident CD4+ and CD8+ T cells confirmed that mCD3-NP facilitated significant increases in targeted LNP uptake and EGFP expression compared with LNP alone **(Fig. 4G, H; Supplementary Fig. 7)**. At a 10-µg total mRNA dose, mCD3-NP addition increased the frequency of DiD+/EGFP+ CD4+ splenic T cells ∼10.5-fold (27% vs 2.5% for uncoated LNPs) and CD8+ T cells ∼19-fold (38.2% vs 2%). This targeted delivery demonstrated a substantial dose-sparing effect; reducing the injected dose to 5 µg maintained robust targeting, yielding a ∼6-fold enhancement in targeted CD4+ T cells (32% vs 5%, respectively) and a ∼12-fold enhancement in targeted CD8+ T cells (35% vs 3%, respectively) relative to untargeted controls.

Similar significant 10- to 20-fold enhancements were also observed for the liver-resident T-cell populations (P<0.0001; **Fig. 4H**). In contrast, while the anti-CD8 NanoPilot resulted in comparatively poor target cell uptake, it still facilitated a significant reduction in hepatic EGFP IVIS signal versus LNP alone (P<0.01; **Fig. 4D, G, H**).

These data suggest that the LDLR-blocking component of NanoPilot is likely the primary driver for reducing hepatic uptake, independent of the targeting ligand’s efficiency. Furthermore, the presence of DiD+/EGFP+ liver-resident T cells in the mCD3-NP group may contribute to the higher residual EGFP IVIS signal observed in the liver. Expression of the activation markers CD69 and CD25 was elevated on both CD4+ and CD8+ T cells in the mCD3-NP, but not mCD8-NP, groups versus controls, while splenic and hepatic T cell counts remained relatively consistent across all treatments **(Supplementary Fig. 8)**. Notably, across all groups AST and ALT enzyme activities remained within the 95% confidence interval of historical reference values,^33^ indicating an absence of acute hepatotoxicity **(Supplementary Fig. 6B)**.

The results from the murine model are therefore consistent with the *in-vitro* assays, supporting the ability of NanoPilot to block off-target uptake and redirect delivery to target cells compared with LNP alone.

### *In-vitro* transfection of human PBMCs

Next we assessed the ability of NanoPilot to deliver payloads to primary human T cells. An initial test with activated and isolated human pan T cells indicated that hCD3-NP led to increased LNP uptake in both primary human CD4+ and CD8+ T cells, with no impact on viable cell counts **(Supplementary Fig. 9A–C)**.

To test whether hCD3-NP was effective at delivering LNPs to unactivated human T cells in peripheral blood mononucleocyte (PBMC) culture, human PBMCs were isolated from buffy coats before transfecting with MC3-DOPE-LP LNPs. The cellular composition of the human PBMC sample was not markedly different after transfection with LNP alone compared with hCD3-NP **(Supplementary Fig. 9D, E)**, indicating that NanoPilot does not impact viability of human immune cells *in vitro*.

Consistent with the previous *in-vitro* and *in-vivo* findings, adding NanoPilot to LNPs resulted in increased on-target (CD4+ and CD8+ cells) and reduced off-target (CD14+ cells) transfection, in both unactivated and activated human PBMCs (**Fig. 5A–C**).

**Figure 5.**
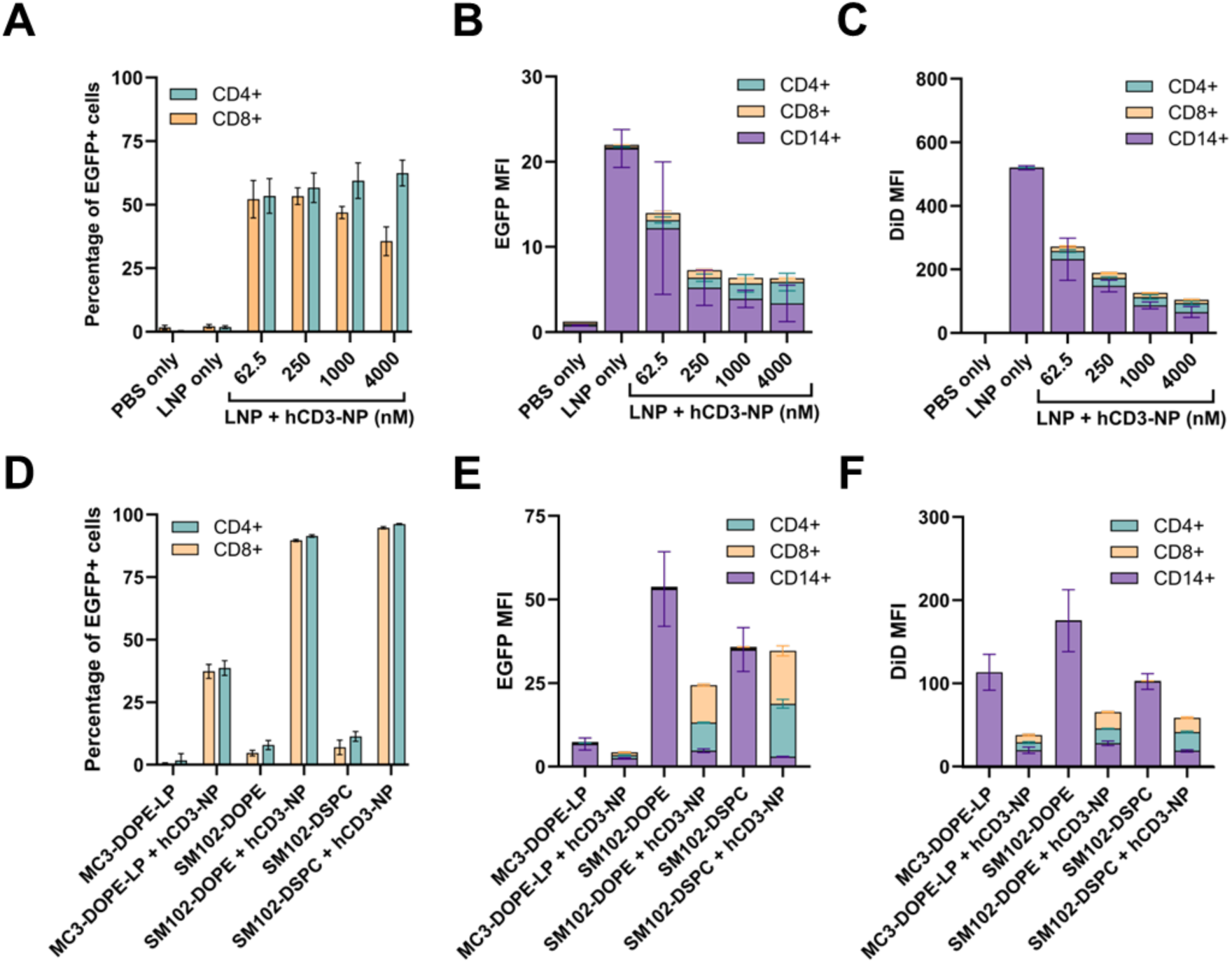
NanoPilot transfects unactivated human T cells within PBMCs. **(A)** Flow cytometric analysis showing hCD3-NP transfection of human pan T cells within unactivated PBMC cultures using MC3-DOPE-LP LNPs. Coating with hCD3-NP increases the transfection (percent EGFP+) of both CD4+ and CD8+ T cells. **(B)** MFI of EGFP and **(C)** DiD following transfection of unactivated human PBMCs. While untargeted MC3-DOPE-LP LNPs exhibit pronounced off-target uptake by monocytes, hCD3-NP titration led to decreased CD14+ uptake and increased CD4+ and CD8+ T cell uptake. **(D)** Comparison of the percentage of EGFP+ cells in unactivated human PBMC mixtures treated with MC3-versus SM102-based LNPs ± 500 nM hCD3-NP. **(E)** EGFP and **(F)** DiD MFI data comparing the targeting profiles of the MC3 and SM-102 formulations ± 500 nM hCD3-NP. EGFP, enhanced green fluorescent protein; hCD3, human CD3; LNP, lipid nanoparticle; MFI, median fluorescence intensity; NP, NanoPilot; PBS, phosphate-buffered saline.

For unmodified LNPs, the CD14+ fraction accounted for nearly all observed LNP uptake and payload translation (**Fig. 5B, C**). Although the addition of NanoPilot (4 μM) achieved an ∼8-fold reduction in this off-target signal, monocyte DiD intensity remained higher than that of target T cells **(Fig. 5C)**. To improve selectivity, we screened alternative ionisable lipids and identified SM-102-based formulations as superior candidates for human T-cell transfection in unactivated PBMCs. Formulation of two SM-102 variants (SM102-DSPC and SM102-DOPE) resulted in homogeneous LNPs (verified via DLS; **Supplementary Table 1; Supplementary Fig. 10)**.

Relative to SM102-DOPE, the SM102-DSPC formulation maximised NanoPilot-mediated delivery to primary T cells while minimising off-target monocyte uptake **(Fig. 5D–F)**. Coating SM102-DSPC LNPs with hCD3-NP resulted in EGFP positivities of 96% and 94% in CD4+ and CD8+ T cells, respectively **(Fig. 5D)**. This corresponded to a ∼40-fold increase in EGFP MFI and a ∼50- to 60-fold increase in DiD MFI compared to untargeted SM102-DSPC LNPs.

Conversely, in off-target CD14+ cells, NanoPilot coating of SM102-DSPC LNPs drove a ∼10-fold decrease in EGFP MFI and a ∼5-fold decrease in DiD MFI **(Fig. 5E, F)**.

## Conclusion

LNP technology has been repeatedly described to hold potential for delivery of novel therapeutics, yet broad applicability has been constrained by off-target uptake and limited cell-specific targeting.^7–9^ The NanoPilot technology described here has demonstrated a dual mechanism *in vivo* and *in vitro*, blocking off-target delivery and redirecting to target cells, thus setting the scene for potential clinical application.

NanoPilot is a modular, easy-to-produce platform designed to redirect LNPs from off-target hepatocytes and immune cells to specific cell types. Its simple structure enhances avidity, comprising an IgG1 scaffold with Fc silencing and LDLR anchors to bind ApoE. The biopharmaceutical characteristics of the backbone are well known, including stability and favourable pharmacokinetics.^34^ Similar to other studies, we found that adjusting the levels of DOPE/DSPC, PEG and SM-102 in the LNP affected ApoE3-mediated uptake and cell-specific targeting.^35–37^

A limitation of the viral vectors currently used in delivery of gene therapies is the long and expensive manufacturing process; for example, current production techniques for CAR-T cell therapies involve *ex-vivo* cell generation, resulting in complex procedures and high costs.^1, 38^ LNP generation is generally quicker, but covalent modification techniques can still take several hours^39, 40^ and introduce chemical moieties that may cause complement activation.^39, 41^ In contrast, the NanoPilot functionalisation process is a rapid, two-step incubation (5 minutes with ApoE, 5 minutes with NanoPilot) that requires no subsequent purification steps, though is amenable to typical cleanup protocols, e.g. tangential flow filtration or dialysis. NanoPilot could therefore be easily scalable for high-throughput production. Our results also demonstrate the modularity of NanoPilot, meaning it can be paired with different antibodies to enable targeting of different cell types. NanoPilot therefore has potential for application in a huge variety of therapy areas.

A particular challenge with LNPs is that formulations that work efficiently in immortalised models (e.g. Jurkat cells) often fail to achieve comparable transfection in primary cells.^37^ Consequently, the ability to rapidly screen different combinations of LNP formulations and targeting domains is crucial for developing targeted therapies. The NanoPilot system directly addresses this need; its highly modular structure facilitates the high-throughput testing necessary to determine the optimal formulation–target pairing for any specific therapeutic application.

Results of the first *in-vitro* and *in-vivo* assays of NanoPilot described here demonstrate an ability to block off-target uptake while enhancing on-target cell delivery and transfection. Other groups have also reported novel approaches to address the current limitations of LNPs. Bimbo et al. combined anti-CD7 and anti-CD3 targeting to create an LNP that codelivered mRNA and DNA to create CAR-T cells *in vivo*; biodistribution data would provide important additional context to their findings.^12^ Hunter et al. adjusted the lipid composition and payload sequence of LNPs in their work on *in-vivo* CAR-T cells; findings indicate desired effects in patient samples, humanised mice and cynomolgus monkeys, alongside residual off-target expression in the tail (mouse model).^11^ Monocyte blocking by NanoPilot reported here compares favourably to a CD47-coated LNP approach.^42^ To the best of our knowledge, we are the first to reduce off-target LNP delivery by competitively blocking the ApoE–LDLR interaction.

Target cell transfection with LNPs can occur alongside off-target delivery;^5^ in a clinical setting, this could reduce functional dose and induce adverse effects. Other groups have reported reduced liver expression with LNPs by inserting certain miRNA sequences into the LNP payload.^43, 44^ However, this does not reduce off-target delivery and the associated risk of side effects. We show that NanoPilot both delivers to and transfects target cells, through measuring DiD and EGFP fluorescence, respectively. Concurrently, NanoPilot blocks both delivery to and transfection of non-target cells.

This research is preliminary, and while we used immune-competent mouse models as well as a murine LDLR-based NanoPilot and murine ApoE in the formulation to maximise relevance, research in additional mouse models is planned. Planned dynamic monitoring of uptake and off-target delivery should provide more information on the off-target blocking currently described at single timepoints. Pending further *in-vitro* and *in-vivo* findings, a Phase 1 clinical trial is needed to assess safety and clinical activity in humans.

In conclusion, the novel NanoPilot technology has demonstrated capability to block and redirect off-target delivery of LNPs, both *in vitro* and *in vivo*. Through this dual mechanism, NanoPilot has the potential to widen the therapeutic application of LNPs. We plan for further work to assess the therapeutic benefits of NanoPilot for multiple indications, and ultimately to investigate tolerability and activity in humans.

## Materials and methods

### NanoPilot cloning

The variable heavy (V_H_) and light (V_L_) chain sequences derived from multiple targeting clones, including the human CD3ε binders cibisatamab and OKT3, the anti-murine clones 2C11 (CD3ε) and YTS-105.18 (CD8α), and the anti-human CD117 clone 9P3, were codon-optimised for expression in Chinese hamster ovary (CHO) cells and produced by gene synthesis (Twist Bioscience). Using Golden Gate DNA assembly these fragments were then inserted into human IgG1 backbones (in a pTwist CMV BG WPRE Neo Vector) to obtain the full-length IgG1 heavy and IgG light chain constructs bearing the complementarity determining regions of interest. The coding region for the human LDLR LA4-LA5 repeat fragment (amino acids V124−A211) and murine LDLR LA4-LA5 fragment (amino acids A124–A212) with an N-terminal rigid linker were similarly gene synthesised and inserted onto the C-terminus of the IgG1 heavy chain via an (EAAAK)_3_ rigid linker to generate the heavy chain of the NanoPilot molecule. For Fc silencing of NanoPilot constructs, variable regions and LDLR fragments were cloned onto an IgG1 backbone sequence synthesised with LALA-PG silencing mutations L234A, L235A and P329G.

### NanoPilot expression

ExpiCHO-S cells were grown to a density of 6×10^6^ cells/mL in ExpiCHO Expression Medium (Gibco) in Optimum Growth Flasks with a 0.2-μm vented cap (Thomson) in a humidified incubator (37°C, 8% CO_2_) with shaking at 120 RPM (19 mm orbit diameter). For transfection, DNA for the IgG1 heavy-LDLR fusions and IgG light chains were complexed at a 1:1 mass ratio using ExpiFectamine CHO reagent and OptiPRO serum-free medium (Gibco) according to the ExpiCHO transfection protocol. After complexation, 1 μg DNA/mL cell culture was added. At 18 hours post-transfection, cells were fed with ExpiCHO feed according to the ExpiCHO protocol and left to express for a further 3–4 days.

### NanoPilot purification

ExpiCHO cell cultures were centrifuged (600g, 10 minutes, 4°C) and the resulting supernatants were clarified through a 0.2-µm PES membrane (Fisherbrand). The clarified media was loaded onto a 5-mL HiTrap Protein G HP column (Cytiva) pre-equilibrated with Protein G binding buffer at 5 mL/min. After washing with 15 column volumes (CV) of binding buffer, the protein was eluted using a 20-CV linear gradient to 100% Protein G elution buffer. Elution fractions were neutralised with 1 M Tris (pH 9.0) at 200 μL/mL of eluate. Pooled fractions were dialysed against DPBS supplemented with 0.5 mM CaCl_2_ and subsequently concentrated to ∼2 mg/mL using a 10-kDa MWCO Amicon Ultra-15 centrifugal device (Millipore). NanoPilots were then sterile filtered using a 0.22-μm Ultrafree^®^ centrifugal filter (Millipore). Final protein concentrations were quantified by A_280_ absorbance. Sample purity and structural homogeneity were verified via SDS-PAGE (4–20% Tris-Glycine) and analytical size-exclusion chromatography (Superdex 200 10/300 GL, Cytiva). See **Supplementary Note** for details on buffer solutions.

### LNP production

Lipid organic phases were prepared by first dissolving lipids in absolute ethanol. The benchmark patisiran-like MC3-DSPC formulation consisted of DLin-MC3-DMA (Cayman Chemical), plant-derived cholesterol (Avanti Research), DSPC (Avanti Research) and DMG-PEG2000 (TebuBio) at a molar ratio of 50:38.5:10:1.5. Variant formulations were prepared by modifying this base ratio: MC3-DOPE utilised DOPE in place of DSPC, whereas the low-PEG variant (MC3-DOPE-LP) utilised DOPE and reduced the DMG-PEG2000 content to 0.75 mol%. Similarly, SM-102 formulations (SM102-DSPC and SM102-DOPE) substituted DLin-MC3-DMA with the SM-102 ionisable lipid (Cayman Chemical) while preserving the 50:38.5:10:1.5 molar ratio of ionisable lipid, cholesterol, helper lipid, and PEG, respectively. To facilitate LNP tracking, 0.25 mol% of DiD (DiIC18(5) solid, Invitrogen) was incorporated into the organic phase prior to mixing.

For *in-vitro* experiments, LNPs were assembled via microfluidic mixing using a NanoAssemblr Spark^TM^ device (Precision NanoSystems). For the aqueous phase, CleanCap EGFP mRNA (TriLink Biotechnologies) was diluted in 100 mM sodium acetate buffer (pH 4.5) to a final concentration of 0.3125 μg/μL. The aqueous phase, the organic phase and RNase-free DPBS dilution buffer were loaded into a NanoAssemblr Spark Cartridge (Cytiva) and microfluidically mixed at a 2:1 (v/v) aqueous:organic ratio.

For *in-vivo* experiments, LNPs were synthesised using a herringbone mixer on a Tamara system (Inside Therapeutics) at an aqueous-to-organic flow rate ratio of 3:1 and a total flow rate of 5 mL/min.

After microfluidic mixing, assembled LNPs were dialysed against 2 L DPBS using Slide-A-Lyzer Dialysis Devices with a 10K MWCO (Thermo Scientific) at 4°C overnight. LNPs were then sterile filtered using a 0.65-μm Ultrafree^®^ centrifugal filter (Millipore) and mRNA concentration and encapsulation efficiency was determined using a modified Quant-it Ribogreen Assay (ThermoFisher Scientific) as described in the GenVoy-ILM User Guide (Precision NanoSystems). Diameter and polydispersity of the LNP formulations used are shown in **Supplementary Table 1**.

### LNP size characterisation

The diameter and concentration of the generated mRNA-LNPs were determined via Nanoparticle Tracking Analysis on a Zetaview PMX-120 (Particle Metrix) using the parameters: measurement type: size; laser wavelength: 488 nm; filter types: scatter or 510 nm (for fluorescence readings); dispersant: PBS; temperature: 21°C; positions measured: 11; 100–200 average counted particles per frame.

### Preparation of NanoPilot-coated LNPs for *in-vitro* titrations

LNPs encapsulating EGFP mRNA were diluted to 2X the desired final dose concentration (typically 20–10 μg/mL) in DPBS+0.5 mM CaCl_2_. The LNP stock was incubated with 2 µg/mL human (Preprotech 350-02-500 μg) or murine (APO-MM102, KACTUS Bio) ApoE3 for 5 minutes at room temperature. NanoPilot was serially diluted in DPBS containing 0.5 mM CaCl_2_ to create a 2X working stock. For transfection, serially diluted NanoPilot was mixed 1:1 with the LNP–ApoE3 mixture and incubated for 5 minutes at room temperature. The LNP–NanoPilot complexes were then added to the cells at a 1:10 LNP:cell culture ratio for a final dose of 100 ng mRNA-LNP per 100,000 cells. Cells were incubated for 18 hours in a humidified incubator (37°C, 5% CO_2_) prior to FACS analysis. Titration concentrations presented in all graphs are the concentration of NanoPilot incubated with the LNP, not the final concentration after addition to cells.

### Jurkat cell culture and transfection

The CD3-(JRT3-T3.5 TCRβ-negative) cell line, which does not express the TCRαβ heterodimer or CD3 on the surface, was obtained from ATCC. The CD3+ JRT3-T3.5 cell line (engineered to express MEL5 TCR and CD8αβ)^45^ was obtained from Lea Knezevic (University of Bristol).

Jurkat cell lines were grown to a density of 0.5–1.5×10^6^ cells/mL in R10 media (RPMI-1640 medium modified with sodium bicarbonate [Sigma], 1X GlutaMAX [Gibco], 10% heat inactivated fetal bovine serum [Gibco], and 10 units/mL penicillin and 100 μg/mL streptomycin) in a humidified incubator (37°C, 5% CO_2_).

To test the effects of NanoPilot on LNP transfection, CD3+ and CD3-Jurkat cells were pelleted by centrifugation (600g, 5 minutes), then washed and resuspended in ImmunoCult-XF T Cell Expansion Medium (STEMCELL Technologies) supplemented with 1 μg/mL human ApoE3 (Merck), 10 units/mL penicillin, 100 μg/mL streptomycin, and 100 μg/mL kanamycin. The cells were mixed at the desired ratio before plating 62,500 cells in a 100-μL volume per well of a flat round-bottomed 96-well TC-Treated Microplate (Corning). The co-culture was then treated with 60 ng/well LNP complexes with or without the hCD3-NP (clone OKT3).

For FACS analysis, Jurkat cells were pelleted in a U-bottomed 96-well TC-Treated Microplate (Corning) plate via centrifugation at 500g. Media were removed and cells resuspended in 50 μL DPBS supplemented with 0.5% w/v BSA and 0.25 μg Human TruStain FcX (BioLegend) to block Fc receptors for 20 minutes at 4°C. For labelling of the CD3+ Jurkat cells, cells were then stained with 50 μL DPBS supplemented with 0.5% w/v BSA and 0.5 μL Human CD8 Alexa Fluor 350 MAb (clone RPA-T8, Bio-Techne) for a further 20 minutes at 4°C. The plate was then centrifuged (500g, 5 minutes) and stain removed before resuspension of the cells in 200 μL PBS + 0.5% w/v BSA. EGFP expression, surface markers and cell viability were determined using flow cytometry on either a BD FACSCanto II with a HTS 96-well plate loader (BD Biosciences) or MACSQuant Analyzer 10 (Miltenyi Biotec). Data were evaluated in FlowJo Software (version 10.1, BD Biosciences).

### ELISA analysis

To test target antigen binding by NanoPilot constructs, indirect ELISAs were performed by first immobilising biotinylated mouse or human CD3εδ heterodimer proteins (CDD-M82W5 and CDD-H82W6, respectively, from ACROBiosystems), or human CD117 (11996-H49H-B Sino Biological) protein at 1 μg/mL in a volume of 100 μL/well of a pre-coated and blocked Streptavidin Microplate (Pierce, 15124). For hCD8-NP testing, murine CD8a/b heterodimer (11146-CD R&D Systems) was immobilised at 1 μg/mL directly to high-bind microplates (Corning 3690). After a 2-hour incubation at room temperature, wells were washed 3 times in 200 μL ELISA wash buffer before applying serial dilutions of NanoPilot constructs for incubation for 1 hour at room temperature. Unbound material was removed by a further 3 washes before applying anti-IgG–HRP conjugate (Mouse Anti-Human IgG Fc-HRP AB99759 from Abcam) for 30-minute incubation at room temperature. For quantification, unbound HRP conjugates were first removed with 6 washes before addition of 100 μL TMB substrate solution (ThermoFisher Scientific) to start the reaction. After a 10-minute incubation at room temperature the reaction was stopped with 100 μL ELISA Stop Solution (SS04, Invitrogen) and colorimetric change determined by measuring absorbance at 460 nm with wavelength subtraction at 570 nm using a plate reader (Cytation 1 Imager, BioTek).

### Murine splenocyte pan T-cell isolation and transfection

Pan T cells were isolated from Balb/C mouse splenocyte suspension (Tebu-bio, IQB-MSP102) by negative selection using a Pan T Cell Isolation Kit (Miltenyi 130-095-130). All steps were performed at 2–8°C. Briefly, splenocytes were resuspended in MACS buffer (PBS pH 7.2, 0.5% BSA, 2 mM EDTA). The cell suspension was incubated with a Biotin-Antibody Cocktail for 5 minutes, followed by a 10-minute incubation with Anti-Biotin MicroBeads (Miltenyi Biotec). The labelled cell suspension was applied to an LS Column (Miltenyi Biotec) within a MACS Separator. The flow-through, containing the enriched, unlabelled pan T cells, was collected. The purity of the isolated T cells was assessed by flow cytometry. All further cell culture was performed in RPMI-1640 medium supplemented with 10% FBS, 1% penicillin-streptomycin, 1% L-glutamine, 40 U/mL murine IL-2 (Miltenyi, 130-120-662) and 0.01 mM 2-mercaptoethanol.

Isolated, unactivated pan T cells were seeded into a 96-well plate at 60,000 cells (100 µL) per well. Diluted LNP–NanoPilot complexes (10 µL) were added to the cells. The plate was incubated overnight (16–20 hours) at 37°C and 5% CO_2_. Post-transfection, cells were analysed for LNP uptake and T-cell subset identification. Cells were first incubated with TruStain FcX™ PLUS (anti-mouse CD16/32, BioLegend) to block Fc receptors. Subsequently, cells were stained with fluorophore-conjugated antibodies against mouse CD4 (PE/Cyanine7, BioLegend, clone RM4-5) and CD8a (APC/Cyanine7, BioLegend, clone 53-6.7). LNP uptake was assessed by measuring the fluorescence of a DiD-labelled lipid within the LNP formulation and translation of the mRNA via EGFP fluorescence. Data were acquired on a MACSQuant Analyser 10 flow cytometer.

### Human PBMC isolation and transfection

Human PBMCs (received from donors from NHS Blood and Transplant) were isolated from buffy coats using the StraightFrom Buffy Coat PBMC Isolation Kit (human; Miltenyi Biotec) according to the manufacturer’s instructions. For transfection assays, isolated unactivated PBMCs were seeded at a density of 65,000 cells/well (100 µL volume) into round-bottom 96-well TC-treated microplates. The cells were cultured in R10 medium supplemented with penicillin/streptomycin and 1 µg/mL human ApoE3. PBMCs were then treated with 65 ng mRNA–LNPs per well (equivalent to 1 µg mRNA per 1×10⁶ cells), formulated either with or without hCD3-NP. Following a 16- to 24-hour incubation (37°C, 5% CO₂), the cells were prepared for flow cytometric analysis to quantify LNP uptake (DiD) and mRNA translation (EGFP). To identify specific immune cell subpopulations, the cells were washed and stained with CD14-VioGreen, CD8-VioBlue and CD4-PE-Vio770 REAfinity antibodies (Miltenyi Biotec) prior to data acquisition.

### Human pan T-cell activation and transfection

Human pan T cells were isolated from RBC-depleted PBMCs by negative selection using the Pan T Cell Isolation Kit (Miltenyi 130-096-535) following the manufacturer’s instructions. Isolated cells were resuspended in ImmunoCult-XF T Cell Expansion Medium (StemCell Technologies) supplemented with 100 IU/mL recombinant human IL-2 (Miltenyi Biotec), 10 units/mL penicillin, and 100 μg/mL streptomycin. Purified T cells were seeded at 1×10^6^ in 24-well flat-bottom plates. Activation was initiated using Human T-Activator CD3/CD28 Dynabeads (Thermo Fisher) at a 1:1 bead-to-cell ratio. Activation was confirmed by flow cytometric analysis of CD69 and CD25 expression. For transfection, activation beads were first removed using a magnetic separator. Cells were seeded at 0.4–0.6×10^6^ cells/mL in the expansion medium supplemented with 1 μg/mL human ApoE3 (Merck). Cells were incubated with 60 ng LNPs for 16–18 hours before analysis of EGFP expression.

### LNP–ApoE blocking assay

An inhibition assay was performed to compare the blocking potential of NanoPilot and the soluble extracellular domain of human LDLR (Ala22–Arg788, LDR-H5224, ACRO Biosystems). The rhLDLR was reconstituted in PBS with 0.5 mM CaCl_2_ to a concentration of 8 µM. The hCD3-NP (OKT3 clone) and rhLDLR were then serially diluted in PBS with 0.5 mM CaCl_2_ from a starting concentration of 4 µM. MC3-DSPC and MC3-DOPE LNP stocks were incubated with human ApoE3 for 5 minutes at room temperature. Subsequently the LNP–ApoE3 complexes were mixed with the serially diluted hCD3-NP or rhLDLR solutions and incubated for 5 minutes. This mixture was then added to CD3-Jurkat T cells for transfection. After 18 hours, GFP median fluorescence intensities were determined via FACS on a MACSQuant Analyser 10 flow cytometer.

### Animal studies

Animal studies were conducted by Crown Bioscience (24-hour C57BL/6 study) and Medicines Discovery Catapult (MDC; 48-hour BALB/c study) in strict accordance with institutional ethical guidelines. Prior to *in-vivo* administration, MC3-DOPE-LP LNP formulations were concentrated to 200 and 100 µg/mL mRNA using 100-kDa MWCO Amicon filters (Millipore) and subsequently sterile-filtered (0.65 µm). Murine ApoE (APO-MM102, KACTUS Bio) was added to the LNPs at concentrations of 40 and 20 µg/mL, respectively. Final dosing solutions were prepared by mixing the concentrated LNPs 1:1 (v/v) with 2x stocks of anti-murine NanoPilot constructs (4 or 2 µM). This yielded final mRNA concentrations of 100 or 50 µg/mL, corresponding to NanoPilot concentrations of 2 or 1 µM, respectively.

For the 24-hour study, 7-week-old female C57BL/6 albino mice (mean body weight 21.4 g; n=5 per group) received a single 100-µL tail vein injection corresponding to a 10 µg or 5 µg total mRNA dose. Control groups received either untargeted LNPs (complexed with murine ApoE only) or a vehicle control (DPBS with 0.5 mM CaCl_2_). A separate 48-hour imaging cohort utilised 7-week-old female BALB/c mice (mean body weight 19 g; n=6 per group).

At the respective 24-hour and 48-hour post-injection timepoints, mice were euthanised and major organs were harvested for *ex-vivo* fluorescence imaging using an IVIS Lumina III *in-vivo* imaging system (PerkinElmer). Fluorescence from the DiD lipid tracer was captured using an excitation filter of 644 nm and an emission filter of 665 nm. EGFP fluorescence from the translated mRNA payload was measured using an excitation filter of 488 nm and an emission filter of 510 nm. For quantitative analysis, regions of interest were drawn around each organ, and the average radiant efficiency ([p/s/cm²/sr]/[µW/cm²]) was calculated using Living Image software (PerkinElmer).

Flow cytometric analysis of murine spleen and liver tissue suspensions was performed 24 hours post-injection using an ID7000C spectral cell analyser (Sony). Following doublet discrimination and dead cell exclusion (Fixable Viability Dye eFluor 780, Invitrogen), viable CD45+ leukocytes were gated into CD4+ and CD8+ T cell subsets. To eliminate EGFP false positives arising from cellular autofluorescence, a sequential gating strategy was employed: T cells were first gated for nanoparticle uptake (DiD+), and the percentage of payload-expressing cells (EGFP+) was subsequently quantified strictly within this DiD+ population. Data were analysed using FlowJo™ v10.10 software (BD Life Sciences).

### Statistical analysis

Data are presented as mean ± standard deviation unless otherwise noted. Statistical significance was determined using an unpaired t-test using GraphPad Prism software. P<0.05 was considered statistically significant.

For the ELISA analysis, curves were fitted with the Agonist vs. response variable slope (four parameter) model within GraphPad Prism (Version 10.2.1) to obtain EC_50_ values.

The half-maximal inhibitory concentration (IC_50_) values for EGFP were calculated by fitting the data to a four-parameter logistic curve using GraphPad Prism.

Targeting ratios were determined by dividing the median EGFP fluorescence intensity (MFI) of the target (e.g., CD3+) cells by the MFI of the non-target (e.g., CD3-) cells.

## Acknowledgements

The authors thank Anne Sayers of Kilns Medical Communications Ltd for providing medical writing support, which was funded by Hone Bio Ltd in accordance with Good Publication Practice guidelines.

The authors gratefully acknowledge NHS Blood and Transplant for providing PBMC materials used in this study. This report is independent research. NHS Blood and Transplant have provided material in support of the research. The views expressed in this publication are those of the author(s) and not necessarily those of NHS Blood and Transplant.

The authors gratefully acknowledge Katy Jepson and Dominic Alibhai of the Wolfson Bioimaging Facility for their support and assistance in this work.

The authors gratefully acknowledge Medicines Discovery Catapult for their *in vivo* research services.

## Reference list

1. Khawar, M.B., et al., Steering the course of CAR T cell therapy with lipid nanoparticles. J Nanobiotechnology, 2024. 22(1): p. 380.

2. Francia, V., et al., The Biomolecular Corona of Lipid Nanoparticles for Gene Therapy. Bioconjug Chem, 2020. 31(9): p. 2046–2059.

3. Bashiri, G., et al., Nanoparticle protein corona: from structure and function to therapeutic targeting. Lab Chip, 2023. 23(6): p. 1432–1466.

4. Gilleron, J., et al., Image-based analysis of lipid nanoparticle-mediated siRNA delivery, intracellular trafficking and endosomal escape. Nat Biotechnol, 2013. 31(7): p. 638–46.

5. Hosseini-Kharat, M., K.E. Bremmell, and C.A. Prestidge, Why do lipid nanoparticles target the liver? Understanding of biodistribution and liver-specific tropism. Mol Ther Methods Clin Dev, 2025. 33(1): p. 101436.

6. Zeng, J., et al., Rapid receptor internalization potentiates CD7-targeted lipid nanoparticles for efficient mRNA delivery to T cells and in vivo CAR T-cell engineering. bioRxiv, 2026.

7. Tenchov, R., et al., Lipid Nanoparticles─From Liposomes to mRNA Vaccine Delivery, a Landscape of Research Diversity and Advancement. ACS Nano, 2021. 15(11): p. 16982–17015.

8. Huang, T., et al., Lipid nanoparticle-based mRNA vaccines in cancers: Current advances and future prospects. Front Immunol, 2022. 13: p. 922301.

9. Larrey, D., et al., Drug-induced liver injury related to gene therapy: A new challenge to be managed. Liver Int, 2024. 44(12): p. 3121–3137.

10. Kazemian, P., et al., Lipid-Nanoparticle-Based Delivery of CRISPR/Cas9 Genome-Editing Components. Mol Pharm, 2022. 19(6): p. 1669–1686.

11. Hunter, T.L., et al., In vivo CAR T cell generation to treat cancer and autoimmune disease. Science, 2025. 388(6753): p. 1311–1317.

12. Bimbo, J.F., et al., T cell-specific non-viral DNA delivery and in vivo CAR-T generation using targeted lipid nanoparticles. J Immunother Cancer, 2025. 13(7).

13. Comirnaty Prescribing Information. February 2026.

14. Patisiran (Onpattro) Prescribing Information. January 2023.

15. Spikevax Prescribing Information. August 2025.

16. Comirnaty Summary of Product Characteristics. April 2026.

17. Patisiran (Onpattro) Summary of Product Characteristics. March 2025.

18. Spikevax Summary of Product Characteristics. February 2026.

19. Wang, J., et al., Recent Advances in Lipid Nanoparticles and Their Safety Concerns for mRNA Delivery. Vaccines (Basel), 2024. 12(10).

20. Bgee. Gene : LDLR - ENSG00000130164 - Homo sapiens (human). [cited 2026 May]; Available from: https://www.bgee.org/gene/ENSG00000130164.

21. Marks, A., et al., mRNA vaccine immunity is enhanced by hepatocyte detargeting and not dependent on dendritic cell expression. Nat Biotechnol, 2026.

22. Park, W., et al., Apolipoprotein Fusion Enables Spontaneous Functionalization of mRNA Lipid Nanoparticles with Antibody for Targeted Cancer Therapy. ACS Nano, 2025. 19(6): p. 6412–6425.

23. Theuerkauf, S.A., et al., ApoE2-DARPin fusion proteins enable selective RNA transfer to CD8 T cells by lipid nanoparticles. J Control Release, 2025. 388(Pt 2): p. 114377.

24. Kayabolen, A., et al., Programmable Lipid Nanoparticle Targeting via Corona Engineering. bioRxiv, 2026.

25. Yan, J., et al., Targeted immunotherapy rescues pulmonary fibrosis by reducing activated fibroblasts and regulating alveolar cell profile. Nat Commun, 2025. 16(1): p. 3748.

26. Rurik, J.G., et al., CAR T cells produced in vivo to treat cardiac injury. Science, 2022. 375(6576): p. 91–96.

27. Bot, A., et al., In vivo chimeric antigen receptor (CAR)-T cell therapy. Nat Rev Drug Discov, 2026. 25(2): p. 116–137.

28. Cevaal, P.M., et al., Efficient mRNA delivery to resting T cells to reverse HIV latency. Nature Communications, 2025. 16(1): p. 4979.

29. Schlothauer, T., et al., Novel human IgG1 and IgG4 Fc-engineered antibodies with completely abolished immune effector functions. Protein Eng Des Sel, 2016. 29(10): p. 457–466.

30. Lo, M., et al., Effector-attenuating Substitutions That Maintain Antibody Stability and Reduce Toxicity in Mice. J Biol Chem, 2017. 292(9): p. 3900–3908.

31. Fisher, C.A. and R.O. Ryan, Lipid binding-induced conformational changes in the N-terminal domain of human apolipoprotein E. J Lipid Res, 1999. 40(1): p. 93–9.

32. Hashiba, K., et al., Balancing Multivalent Avidity and Receptor Availability Governs mRNA Delivery by Antibody-Functionalized Lipid Nanoparticles. Nano Lett, 2026. 26(14): p. 4630–4641.

33. Charles River. C57BL/6 Mouse Hematology and Biochemistry. [cited 2026 May]; Available from: https://www.criver.com/products-services/find-model/c57bl6-mouse?region=3671.

34. Lobner, E., et al., Engineered IgG1-Fc--one fragment to bind them all. Immunol Rev, 2016. 270(1): p. 113–31.

35. Zhang, R., et al., Helper lipid structure influences protein adsorption and delivery of lipid nanoparticles to spleen and liver. Biomater Sci, 2021. 9(4): p. 1449–1463.

36. Kim, M., et al., Engineered ionizable lipid nanoparticles for targeted delivery of RNA therapeutics into different types of cells in the liver. Sci Adv, 2021. 7(9).

37. VanKeulen-Miller, R., et al., Customizable mRNA Lipid Nanoparticles for Transfection of Primary Human T Cells. ACS Nano, 2025. 19(49): p. 41836–41849.

38. Kong, Y., et al., CAR-T cell therapy: developments, challenges and expanded applications from cancer to autoimmunity. Front Immunol, 2024. 15: p. 1519671.

39. Zaleski, M.H., et al., Conjugation Chemistry Markedly Impacts Toxicity and Biodistribution of Targeted Nanoparticles, Mediated by Complement Activation. Adv Mater, 2025. 37(5): p. e2409945.

40. Chen, M.Z., et al., A versatile antibody capture system drives specific in vivo delivery of mRNA-loaded lipid nanoparticles. Nat Nanotechnol, 2025.

41. Geisler, H.C., et al., Preparation of targeted lipid nanoparticles for precision nucleic acid delivery. Nat Protoc, 2026.

42. Papp, T.E., et al., CD47 peptide-cloaked lipid nanoparticles promote cell-specific mRNA delivery. Mol Ther, 2025. 33(7): p. 3195–3208.

43. Parrett, B.J., S. Yamaoka, and M.A. Barry, Reducing off-target expression of mRNA therapeutics and vaccines in the liver with microRNA binding sites. Mol Ther Methods Clin Dev, 2025. 33(1): p. 101402.

44. Xu, S., et al., In vivo genome editing of human haematopoietic stem cells for treatment of blood disorders using mRNA delivery. Nat Biomed Eng, 2025.

45. Clement, M., et al., CD8 coreceptor-mediated focusing can reorder the agonist hierarchy of peptide ligands recognized via the T cell receptor. Proc Natl Acad Sci U S A, 2021. 118(29).

