## Supplementary Material for "Extrahepatic, cell-specific delivery of LNPs through competitive inhibition of ApoE-mediated uptake"

### Supplement

#### Supplementary Notes

HepG2 monoculture and HepG2-Kasumi-1 co-culture assays

Jurkat T cell imaging

Additional methodological details

#### Supplementary Tables

**Supplementary table 1.** Dynamic light scattering data showing mean diameter and polydispersity of the LNP formulations listed in this study.

**Supplementary table 2.** IC50 values determined for rhLDLR- and hCD3-NP-mediated inhibition of ApoE3-mediated EGFP-LNP expression in CD3- cells.

#### Supplementary Figures

**Supplementary figure 1.** Purification and ELISA data of the NanoPilot construct.

**Supplementary figure 2.** Dynamic light scattering data of the MC3 LNP formulations used in this study and the MC3-DOPE-LP formulation with and without ApoE3 and hCD3-NP.

**Supplementary figure 3.** Comparison of anti-CD3ε antibody clones for NanoPilot constructs.

**Supplementary figure 4.** Optimisation of LNP lipid formulation enhances NanoPilot targeting efficiency.

**Supplementary figure 5.** Flow cytometric analysis of unactivated murine pan T cells transfected with MC3-DOPE-LP with and without mCD3-NP and mCD8-NP.

**Supplementary figure 6.** *Ex-vivo* analysis of livers from LNP-dosed mice.

**Supplementary figure 7.** Flow cytometry gating strategy for the identification of DiD+/EGFP+ T cells.

**Supplementary figure 8.** NanoPilot LNP delivery impacts murine T-cell activation in vivo without severely depleting splenic or intrahepatic populations.

**Supplementary figure 9.** Flow cytometry analysis of transfected human primary PBMCs.

**Supplementary figure 10.** Dynamic light scattering data of the SM102-DOPE and SM102-DSPC formulations alongside the MC3-DOPE-LP formulation as comparison.

#### Supplementary Notes

*HepG2 monoculture and HepG2-Kasumi-1 co-culture assays*

The human hepatocellular carcinoma cell line HepG2 (ATCC HB-8065) was cultured in R10 medium. Cells were maintained in T-75 flasks in a humidified incubator (37°C with 5% $CO_2$). The medium was replaced every 2–3 days. Cells were sub-cultured upon reaching 70% confluency. The monolayer was washed with PBS and detached using TrypLE Express trypsin enzyme (Gibco). The trypsin was neutralised with R10 medium, and the cell suspension was centrifuged (100g, 5 minutes). The cell pellet was resuspended in fresh medium, and viable cell concentration was determined by Trypan Blue exclusion.

For transfection assays, HepG2 cells were re-seeded at a density of 45,000 cells/well in flat-bottomed TC treated microplates (Corning) and incubated for 24 hours prior to transfection. For the HepG2 monoculture inhibition experiments, MC3-DSPC LNPs (10 µg/mL mRNA) were coated with 2 µg/mL ApoE3 prior to incubation with a titration range of hCD3-NP. The HepG2 cells were treated with 100 ng mRNA per well of the resulting LNP complexes, or untargeted controls. Following an 18-hour incubation, the HepG2 cells were detached using TrypLE Express and EGFP expression was quantified on a BD FACSCanto II flow cytometer.

For the hCD117-NP targeting assay utilising a co-culture model, Kasumi-1 cells (ATCC-CRL-2724; maintained in R10 medium and split every 3–5 days) were pre-stained with CellTrace™ Far Red (Invitrogen). The medium was removed from adherent HepG2 cells, and 35,000 pre-stained Kasumi-1 cells were plated on top in 100 µL of ImmunoCult-XF T Cell Expansion Medium (supplemented with 1 µg/mL human ApoE3, 1% penicillin-streptomycin, and 100 µg/mL kanamycin). The co-culture was treated with 100 ng mRNA per well of MC3-DSPC LNPs formulated with or without hCD117-NP (clone 9P3). Following a 16-hour incubation, the mixed cell population was detached using TrypLE Express and EGFP expression was quantified on a BD FACSCanto II flow cytometer.

*Jurkat T cell imaging*

Jurkat cells (CD3+ and CD3−) were stained with 5 µg/mL Hoechst 33342 (Tocris Bioscience) for 10 min at room temperature. Cells were seeded at 65,000 cells/well into fibronectin-coated (12.5 µg/mL) 96-well optical-bottom plates (Thermo Scientific) in 100 µL R10 medium supplemented with 1 µg/mL human ApoE3 (Merck). Cells were transfected in triplicate with 60 ng MC3-DOPE-LP LNPs formulated with or without hCD3-NP. Plates were incubated at 37°C with 5% $CO_2$, and imaging commenced within 2 hours. Live-cell imaging was performed using a Cell Discoverer 7 widefield microscope (Carl Zeiss) with a 20x/0.7 NA objective (Wolfson Bioimaging Facility, University of Bristol). Three regions per well were imaged hourly for 24 hours across EGFP, DiD, Hoechst, and brightfield channels. Images were processed in Fiji using the Modular Image Analysis (MIA) plugin.[1,2] Cell detection and temporal tracking were performed on extended-focus brightfield images using Cellpose.[3] Fluorescence intensities were quantified from 3D maximum intensity projections and corrected for photobleaching using the simple ratio method.[4] Single-cell median EGFP intensities were pooled, and the 80th percentile of the raw fluorescence was calculated per condition. Background and signal drift were corrected by subtracting the time-matched 80th-percentile values of the PBS controls. Cumulative EGFP expression was quantified using area under the curve (AUC) analysis on these background-subtracted trajectories in GraphPad Prism (Version 10.2.1).

*Additional methodological details*

Protein G binding buffer: 25 mM $NaPO_4$ pH 7.0, 150 mM NaCl, 0.5 mM $CaCl_2$

Protein G elution buffer: 100 mM citric acid pH 2.5, 150 mM NaCl, 0.5 mM $CaCl_2$

IEX-A buffer: 50 mM TRIS pH 8.0, 0.5 mM $CaCl_2$
IEX-B: 50 mM TRIS pH 8.0, 30 mM NaCl, 0.5 mM $CaCl_2$
ELISA wash buffer: 130 mM NaCl, 20 mM TRIS pH 7.5, 0.05% Tween-20, 0.1% BSA

*References for supplementary notes*


1. Schindelin, J., et al., *Fiji: an open-source platform for biological-image analysis.* Nat Methods, 2012. **9**(7): p. 676-82.
2. Cross, S.J., J. Fisher, and M.A. Jepson, *ModularImageAnalysis (MIA): Assembly of modularised image and object analysis workflows in ImageJ.* J Microsc, 2024. **296**(3): p. 173-183.
3. Stringer, C., et al., *Cellpose: a generalist algorithm for cellular segmentation.* Nat Methods, 2021. **18**(1): p. 100-106.
4. Miura, K., *Bleach correction ImageJ plugin for compensating the photobleaching of time-lapse sequences.* F1000Res, 2020. **9**: p. 1494.

#### Supplementary tables

| LNP formulation | Mean effective diameter (nm) | Mean polydispersity |
|---|---|---|
| MC3-DSPC | 97.1 | 0.184 |
| MC3-DOPE | 115.8 | 0.143 |
| SM102-DSPC | 95.5 | 0.062 |
| SM102-DOPE | 103.2 | 0.117 |
| MC3-DOPE-LP | 98.2 | 0.140 |
| MC3-DOPE-LP + ApoE3 | 101.0 | 0.149 |
| MC3-DOPE-LP + ApoE3 + hCD3-NP | 102.1 | 0.141 |

**Supplementary table 1.** Dynamic light scattering data showing mean diameter and polydispersity of the LNP formulations listed in this study.
ApoE3, apolipoprotein E3; hCD3, human CD3; LNP, lipid nanoparticle; NP, NanoPilot.

| Sample | MC3-DOPE LNP $IC_{50}$ (nM) | MC3-DSPC LNP $IC_{50}$ (nM) |
|---|---|---|
| hCD3-NP | 6.04 (95% CI: 5.490-6.652) | 2.43 (95% CI: 1.657-3.430) |
| rhLDLR | 49.33 (95% CI: 42.37-57.63) | 1.75 (95% CI: 1.159-2.503) |

**Supplementary table 2.** $IC_{50}$ values determined for rhLDLR- and hCD3-NP-mediated inhibition of ApoE3-mediated EGFP-LNP expression in CD3- cells. Both LNPs are 1.5% PEG patisiran-like formulations. Curves were fitted with the agonist vs. response variable slope (four parameter) model within GraphPad Prism (Version 10.2.1).
ApoE3, apolipoprotein E3; CI, confidence interval; EGFP, enhanced green fluorescent protein; hCD3, human CD3; $IC_{50}$, half maximal inhibitory concentration; LNP, lipid nanoparticle; NP, NanoPilot; rhLDLR, recombinant human low density lipoprotein receptor.

#### Supplementary figures

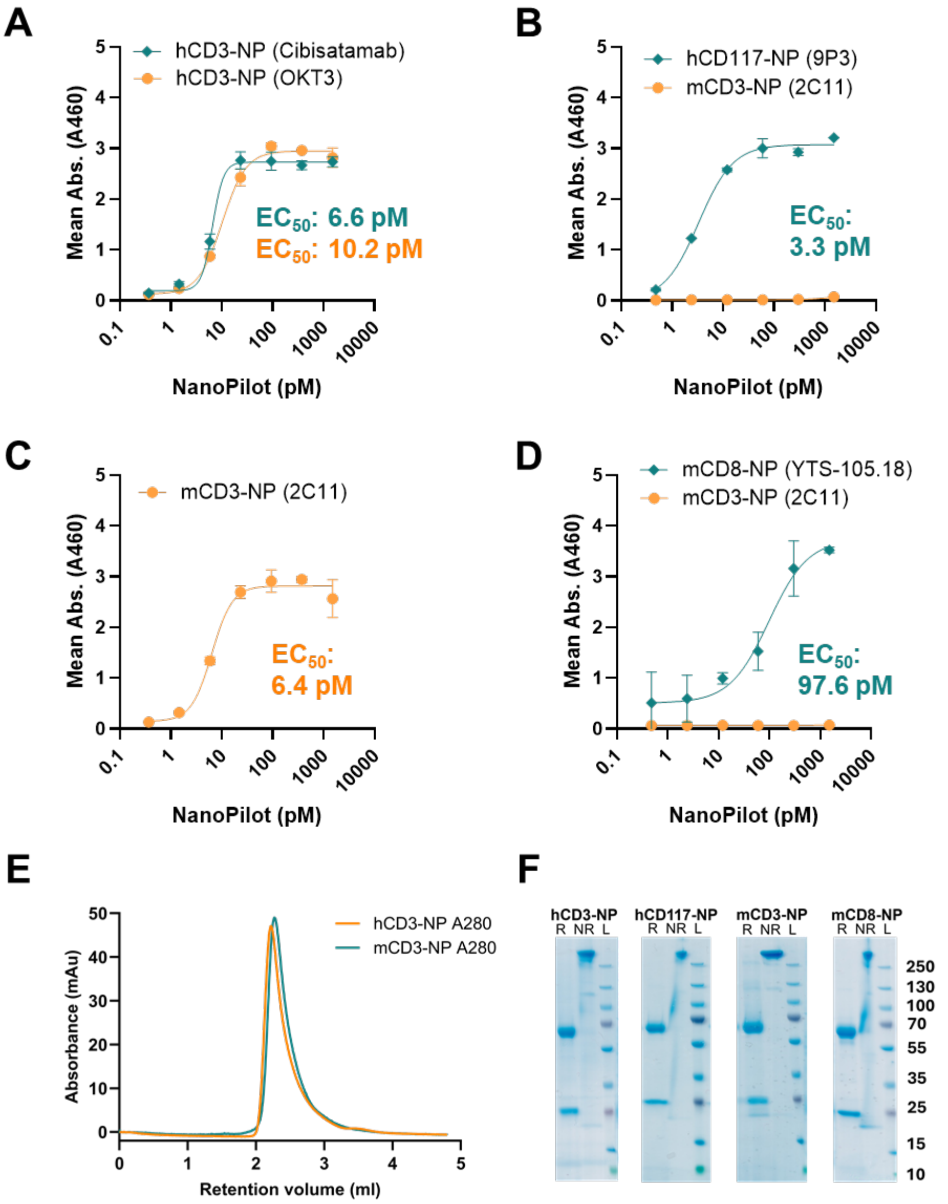


**Supplementary figure 1. Purification and ELISA data of the NanoPilot construct. (A)** ELISA data showing binding of hCD3-NP (cibisatamab and OKT3 variable region variants) to biotinylated human CD3 immobilised on a streptavidin plate. **(B)** hCD117-NP and negative control mCD3-NP binding to biotinylated human CD117 immobilised on streptavidin plate. **(C)** mCD3-NP binding to biotinylated murine CD3ε immobilised on streptavidin plate. **(D)** mCD8-NP and negative control mCD3-NP binding to murine CD8 alpha/beta heterodimer immobilised directly to a high binding plate. **(E)** Representative size exclusion profiles of purified NanoPilot material showing homogeneity and lack of aggregates (run on Superose 6 increase 5/150 GL). **(F)** SDS-PAGE of purified NanoPilot constructs in reducing (R) and non-reducing (NR) loading dye.
Calculated $EC_{50}$ values are depicted for each ELISA curve. Values were determined by fitting an agonist vs response variable slope (four parameter) model within GraphPad Prism (Version 10.2.1).

$EC_{50}$, half maximal effective concentration; ELISA, enzyme-linked immunosorbent assay; hCD117/3, human CD117/3; mCD3/8, murine CD3/8; NP, NanoPilot; NR, non-reducing; R, reducing; SDS-PAGE, sodium dodecyl sulfate polyacrylamide gel electrophoresis.

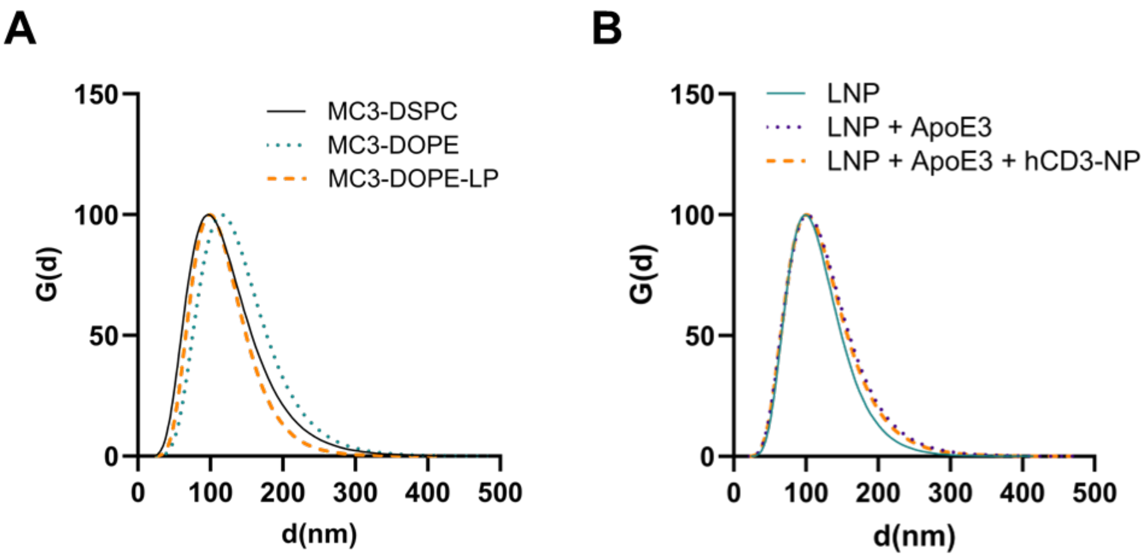


**Supplementary figure 2.** Dynamic light scattering data of **(A)** the MC3 LNP formulations used in this study and **(B)** the MC3-DOPE-LP formulation with and without ApoE3 and hCD3-NP. All curves shown represent intensity-weighted lognormal data.

ApoE3, apolipoprotein E3; hCD3, human CD3; LNP, lipid nanoparticle; NP, NanoPilot.

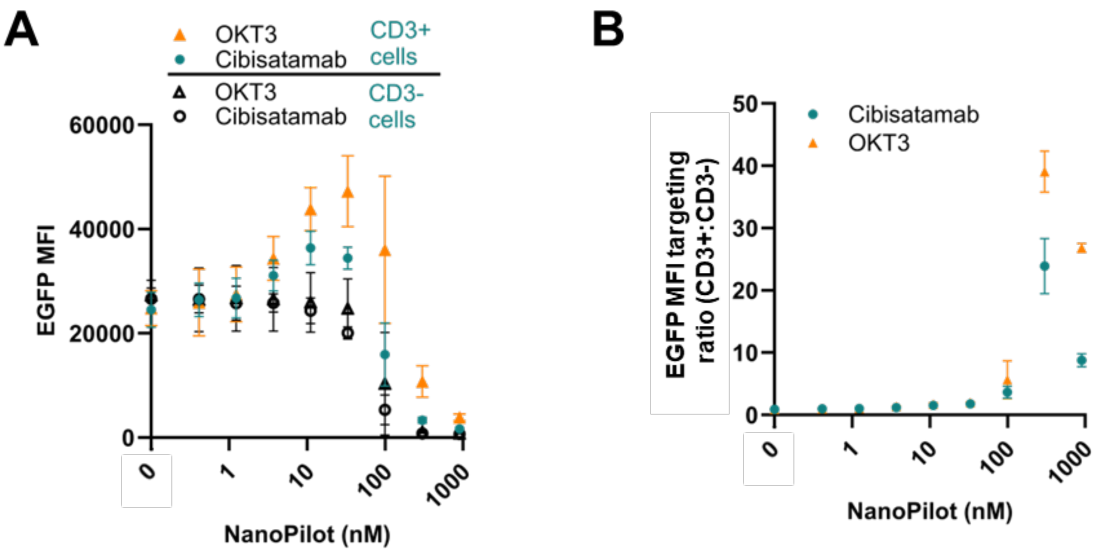


**Supplementary figure 3. Comparison of anti-CD3ε antibody clones for NanoPilot constructs. (A)** Flow cytometric comparison of CD3+ targeted MC3-DSPC LNP delivery in CD3+/CD3- Jurkat cell co-cultures utilising cibisatamab- vs OKT3-derived NanoPilot constructs. **(B)** EGFP median targeting ratio of OKT3-derived NanoPilot demonstrated superior targeted transfection efficiency versus cibisatamab-based NanoPilot.

Data are presented as mean ± SD (n=3). EGFP, enhanced green fluorescent protein; hCD3, human CD3; MFI, median fluorescence intensity; NP, NanoPilot; SD, standard deviation.

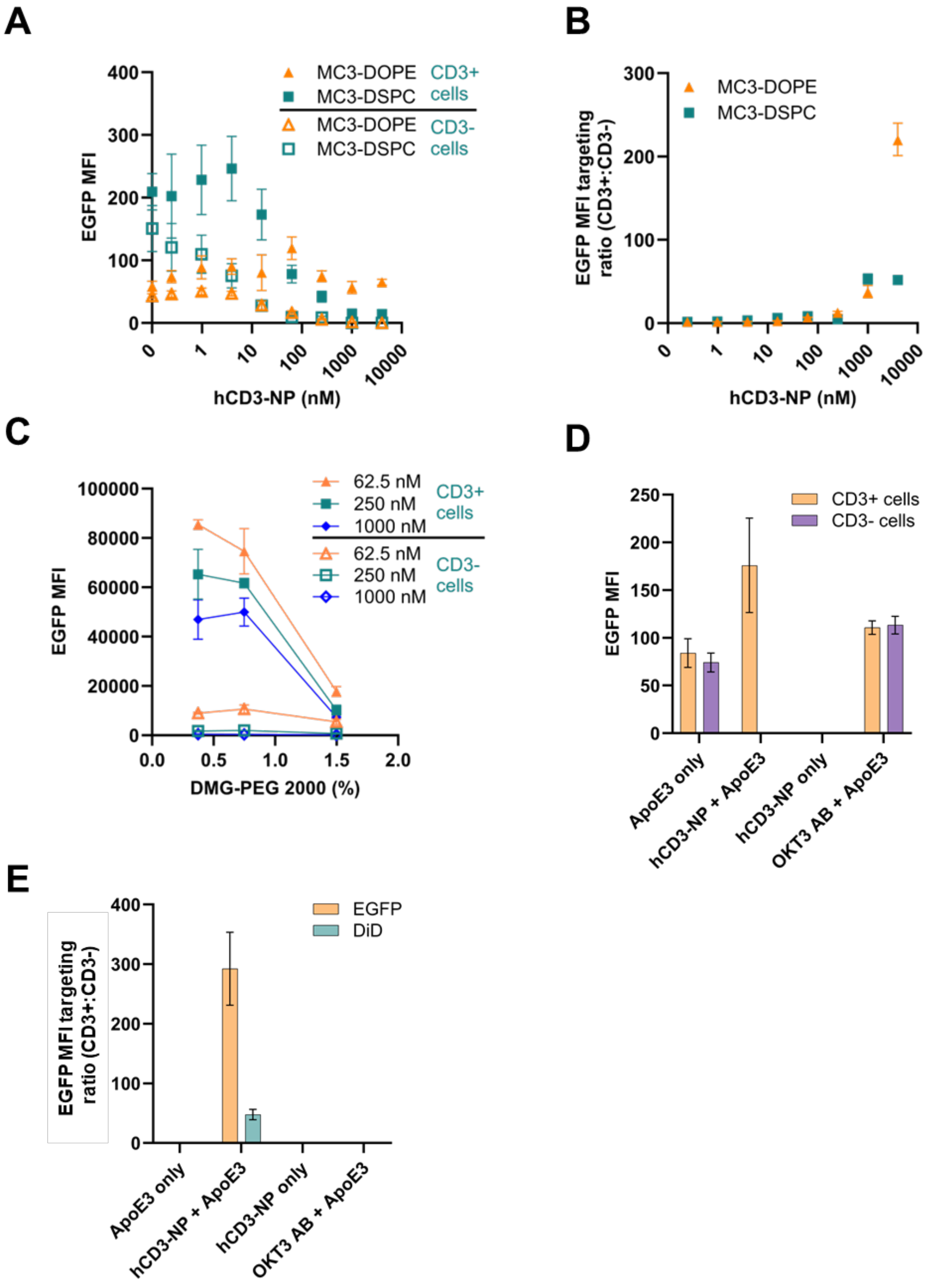


**Supplementary figure 4. Optimisation of LNP lipid formulation enhances NanoPilot targeting efficiency. (A)** Flow cytometry data showing EGFP MFI in target (CD3+) and off-target (CD3-) Jurkat T cells (co-culture model) treated with hCD3-NP complexed with MC3-DOPE vs MC3-DSPC. **(B)** Comparison of the targeting ratio for MC3-DSPC and MC3-DOPE LNP formulations across a NanoPilot titration. **(C)** EGFP expression in CD3+/CD3- cells treated with the MC3-DOPE-LNP formulations at varying concentrations, containing 1.5% (MC3-DOPE), 0.75% (MC3-DOPE-LP) or 0.375% DMG-PEG2000. **(D)** Flow cytometric quantification of MFI and **(E)** targeting ratios in a CD3+/CD3- Jurkat cell co-culture treated with MC3-DOPE-LP LNPs (formulated with 0.25 mol% DiD), complexed with 500 nM of either hCD3-NP or an unmodified OKT3 control antibody ± ApoE3. The targeted uptake of LNPs is completely abolished upon the removal of ApoE3 or when utilising a control OKT3 antibody lacking the LDLR-binding fragment.

Data are presented as mean ± SD (n=3). AB, antibody; ApoE3, apolipoprotein E3; EGFP, enhanced green fluorescent protein; hCD3, human CD3; LDLR, low density lipoprotein receptor; LNP, lipid nanoparticle; MFI, median fluorescence intensity; NP, NanoPilot.

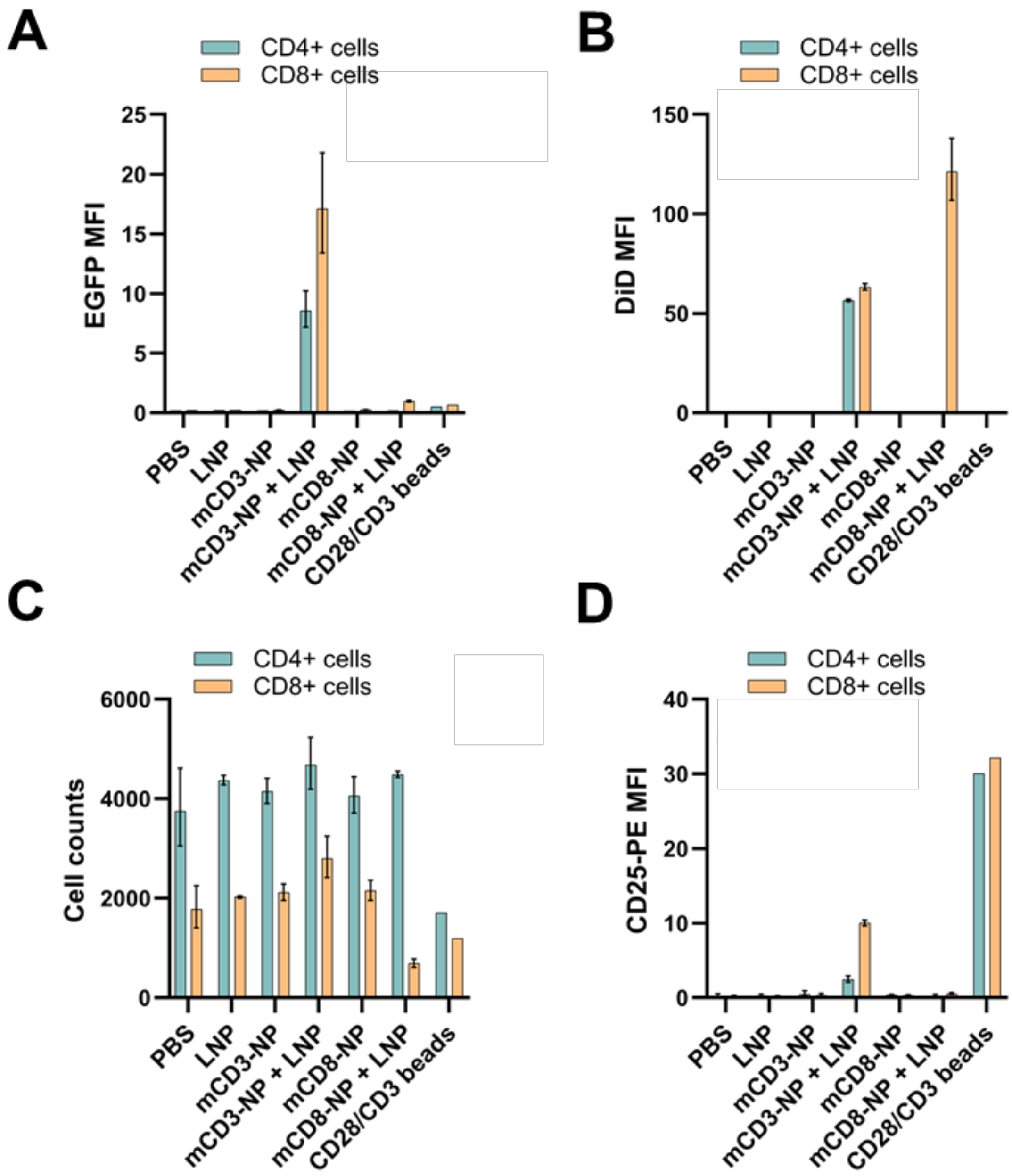


**Supplementary figure 5. Flow cytometric analysis of unactivated murine pan T cells transfected with MC3-DOPE-LP with and without mCD3-NP and mCD8-NP. (A)** EGFP MFI data show considerably higher EGFP expression in CD4+ and CD8+ T cells with mCD3-NP versus the other conditions. **(B)** Conversely, DiD MFI values are higher for mCD8-NP than mCD3-NP, specifically for CD8+ T cells. **(C)** Stable cell counts in LNP and LNP+NanoPilot conditions. **(D)** Late activation marker CD25 MFI data with CD28/CD3 activation bead-positive control showing higher activation for mCD3-NP and no activation for mCD8-NP.
Data are presented as mean ± SD (n=3). EGFP, enhanced green fluorescent protein; LNP, lipid nanoparticle; mCD3/8, murine CD3/8; MFI, median fluorescence intensity; NP, NanoPilot; PBS, phosphate-buffered saline.

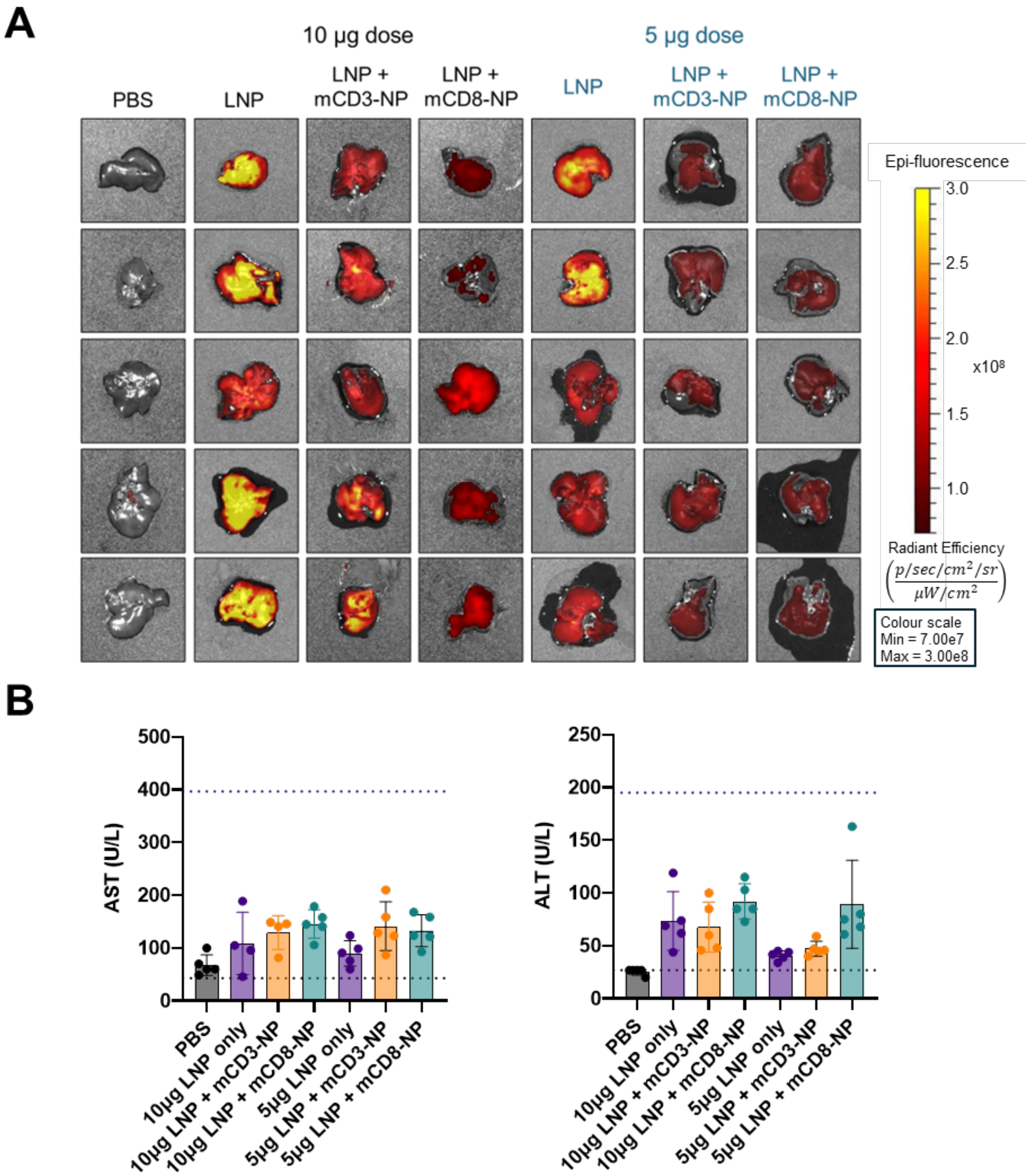


**Supplementary figure 6. *Ex-vivo* analysis of livers from LNP-dosed mice. (A)** Raw *ex-vivo* IVIS EGFP images of C57BL/6 mouse livers 24 hours post-injection of MC3-DOPE-LP ± NanoPilot alongside PBS controls. **(B)** AST (left) and ALT (right) enzyme concentrations measured from murine serum. Dotted lines represent low and high reference ranges for ALT (27–195 U/L) and AST (43–397 U/L) (Charles River. C57BL/6 Mouse Hematology and Biochemistry. https://www.criver.com/products-services/find-model/c57bl6-mouse?region=3671. Accessed May 2026).

ALT, alanine aminotransferase; AST, aspartate aminotransferase; EGFP, enhanced green fluorescent protein; IVIS, *in vivo* imaging system; LNP, lipid nanoparticle; mCD3/8, murine CD3/8; NP, NanoPilot; PBS, phosphate-buffered saline.

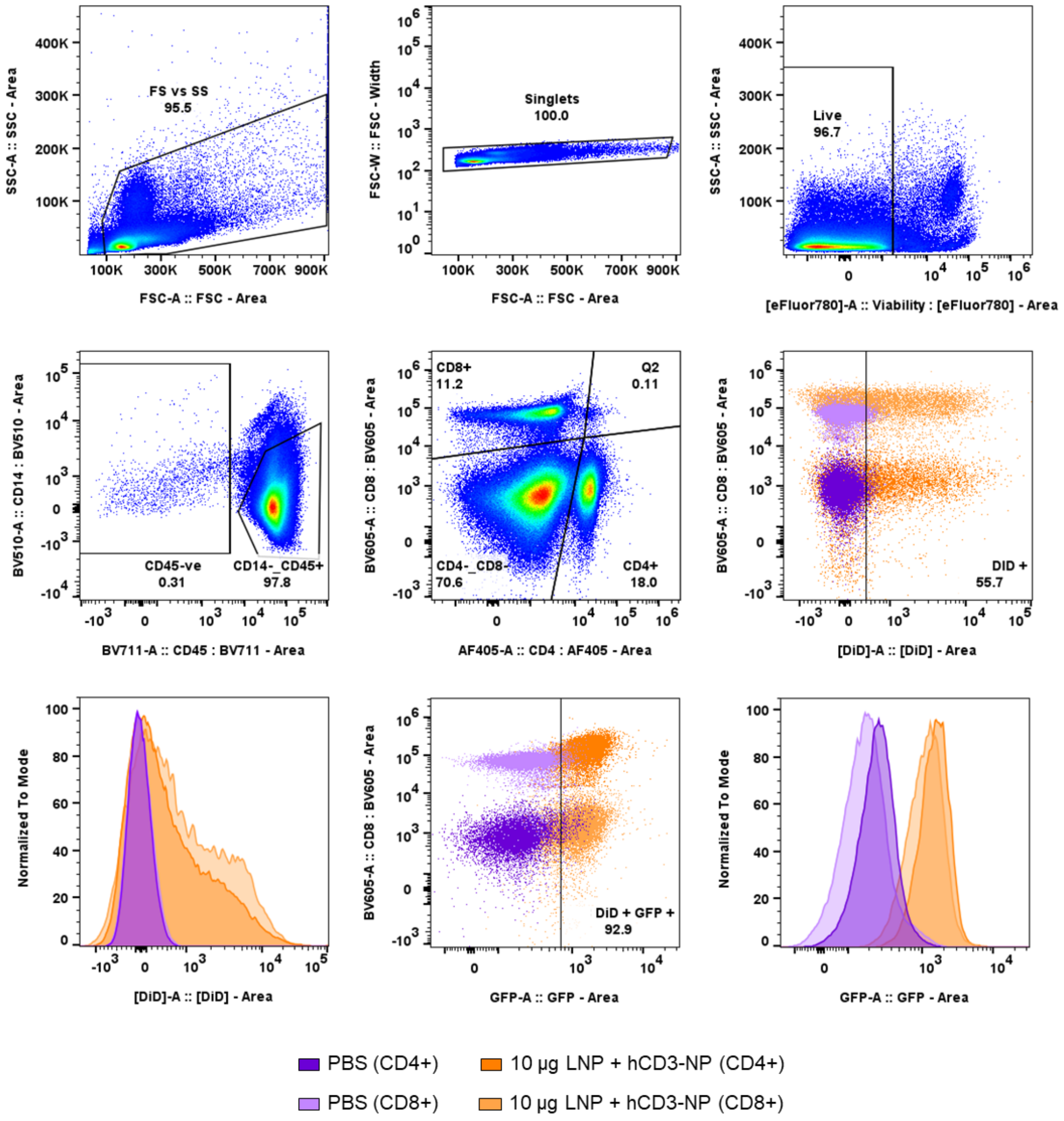


**Supplementary figure 7. Flow cytometry gating strategy for the identification of DiD+/EGFP+ T cells.** Representative plots from mouse spleen samples taken 24 hours post-injection. EGFP, enhanced green fluorescent protein; FCS, forward scatter; hCD3, human CD3; LNP, lipid nanoparticle; NP, NanoPilot; PBS, phosphate-buffered saline; SSC, side scatter.

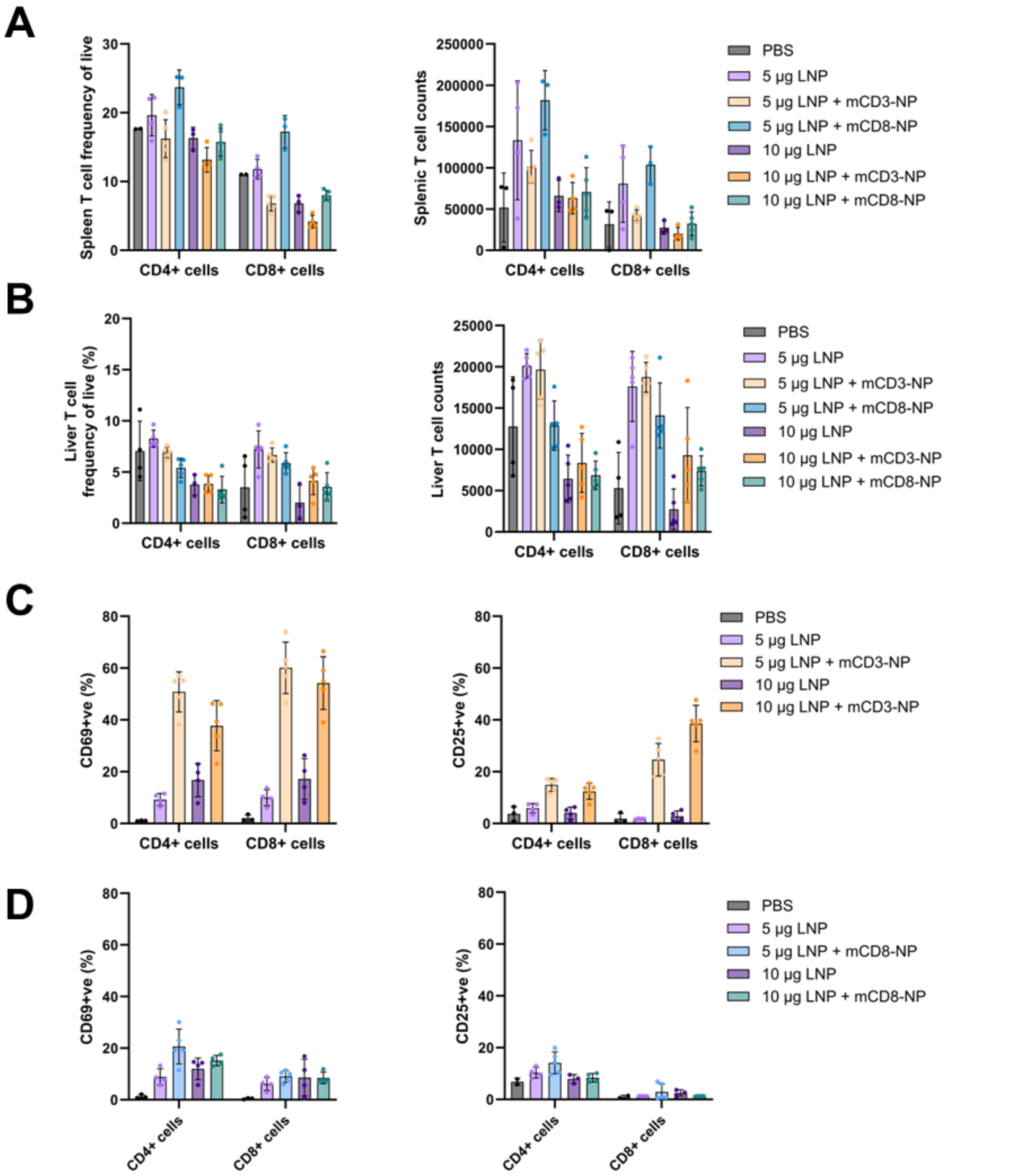


**Supplementary figure 8**. **NanoPilot LNP delivery impacts murine T-cell activation *in vivo* without severely depleting splenic or intrahepatic populations. (A, B)** Flow cytometric quantification of CD4+ and CD8+ T cell maintenance in **(A)** the spleen and **(B)** liver following systemic administration of NanoPilot LNPs. Data show the frequency of live cells (%) and absolute cell counts across PBS, untargeted LNP, and targeted (mCD3-NP or mCD8-NP) LNP groups at both 5-µg and 10-µg total mRNA doses. **(C)** Targeted delivery via mCD3-NP induces *in-vivo* activation of both CD4+ and CD8+ splenic T cells, quantified by the upregulation of early (CD69) and late (CD25) activation markers compared to untargeted controls. **(D)** Evaluation of CD69 and CD25 upregulation on splenic CD4+ and CD8+ T cell subsets following administration of mCD8-NP targeted LNPs.

Data are presented as mean ± SD (n=5 biologically independent animals per group, except for the splenic PBS [n=3], liver PBS [n=4] and splenic 10 µg untargeted LNP [n=4] controls due to sample loss during *ex-vivo* tissue processing).

mCD3/8, murine CD3/8; LNP, lipid nanoparticle; NP, NanoPilot; PBS, phosphate-buffered saline; SD, standard deviation.

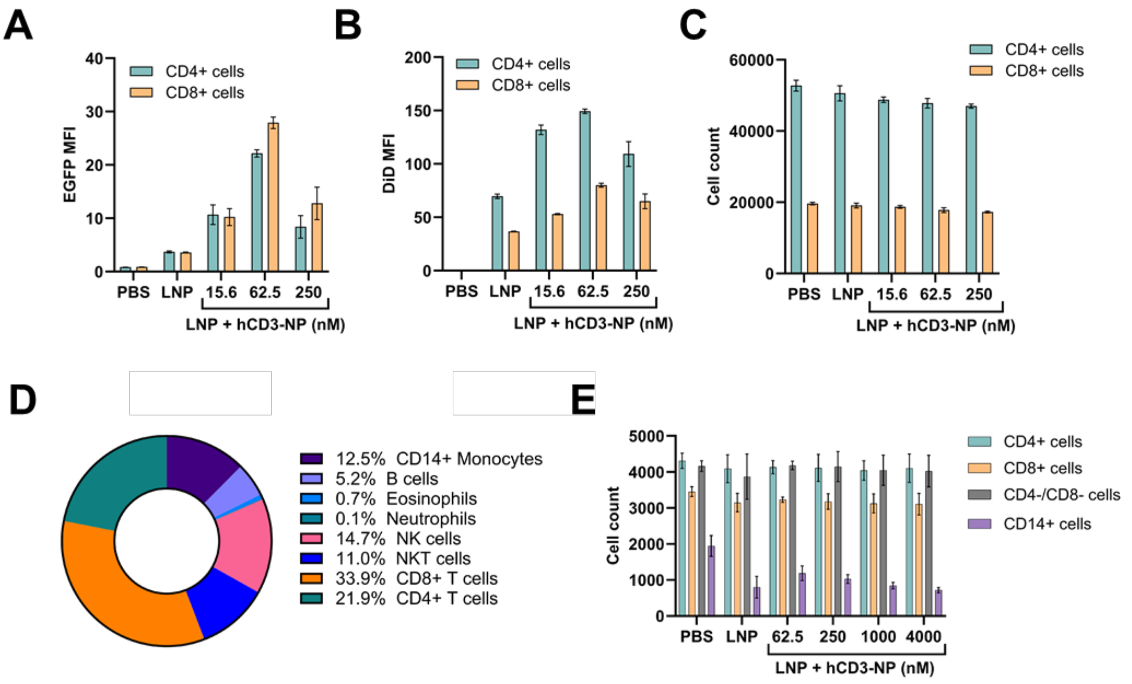


**Supplementary figure 9. Flow cytometry analysis of transfected human primary PBMCs. (A)** Flow cytometry median EGFP and **(B)** DiD fluorescence intensity data for activated human pan T cells with MC3-DOPE-LP LNPs ± hCD3-NP. **(C)** Viable cell count data for the same pan T cell experiment showing no reduction in viability across the NanoPilot titration range. **(D)** Pre-transfection cellular composition of unactivated human PBMC samples isolated from Buffy coat (FACS analysis). **(E)** Post-transfection PBMC cell counts showing incubation with NanoPilot does not affect the distribution of cell types versus controls. EGFP, enhanced green fluorescent protein; FACS, fluorescence-activated cell sorting; hCD3, human CD3; LNP, lipid nanoparticle; MFI, median fluorescence intensity; NK, natural killer; NP, NanoPilot; PBMC, peripheral blood mononucleocyte; PBS, phosphate-buffered saline.

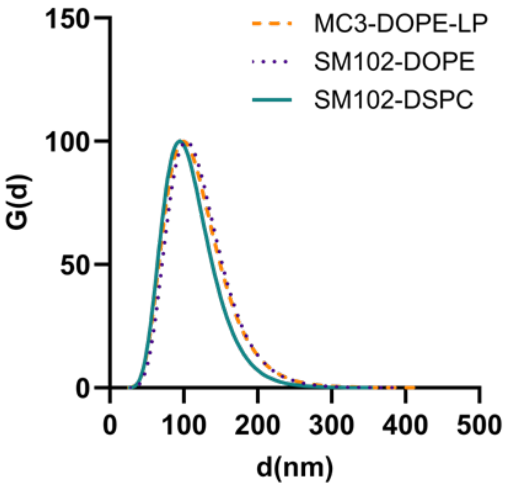


**Supplementary figure 10.** Dynamic light scattering data of the SM102-DOPE and SM102-DSPC formulations alongside the MC3-DOPE-LP formulation as comparison. All curves shown represent intensity-weighted lognormal data.
